# Multi-omic analyses unveil contrasting composition and spatial distribution of specialized metabolites in seeds of *Camelina sativa* and other Brassicaceae

**DOI:** 10.1101/2024.05.31.596893

**Authors:** Léa Barreda, Céline Brosse, Stéphanie Boutet, Nicolas Klewko, Delphine De Vos, Tracy Francois, Boris Collet, Damaris Grain, Céline Boulard, Jean Chrisologue Totozafy, Benoît Bernay, François Perreau, Loïc Lepiniec, Loïc Rajjou, Massimiliano Corso

## Abstract

Seeds of Brassicaceae produce a large diversity of beneficial and antinutritional specialized metabolites (SMs) that influence their quality and provide resistance to stresses. While the distribution of these compounds has been described in leaves and roots tissues, limited information is available about their spatio-temporal accumulation in seeds.

*Camelina sativa* (camelina) is an oilseed Brassicaceae cultivated for human and animal nutrition, and for industrial uses. While we previously explored SM diversity and plasticity, no information is available about SM distribution and expression of related proteins and genes in camelina seeds.

In this study, we used a multi-omic approach, integrating untargeted metabolomics, data-independent acquisition proteomics, and transcriptomics to investigate the synthesis, modifications and degradations of SMs accumulated in the different seed tissues (i.e. seed coat, endosperm, and embryo) at 6 developmental and 2 germination stages. Our results showed distinct patterns of SMs and their related pathways, highlighting significant contrasts in seed composition and spatial distribution for the defence-related and antinutritional glucosinolate (GSL) compounds among camelina, *Arabidopsis thaliana,* and *Brassica napus,* three closely-related Brassicaceae species. Notably, the variation in GSL spatial distributions was primarily driven by differences in their structure and transport mechanisms. Long chain C8-C11 methylsulfinylalkyl GSLs were predominantly accumulated in the seed coat and endosperm, while mid- and short-chain C3-C7 methylsulfinylalkyl GSLs were accumulated in the embryo.

Characterizing the spatial dynamics of seed SMs provides valuable insights that can guide the development of crops with optimized distribution of beneficial and toxic metabolites, improving seed nutritional profiles for feed and food.

## Introduction

Plants produce a myriad of specialized metabolites (SMs) with tremendous interest for agriculture, nutrition and health (Arimura and Maffei, 2016). The large diversity of chemical structures and, hence, of functions observed among specialized metabolic classes arises from different functional groups (decorations or modifications) commonly added to SM core structures (Barreda et al., 2024). SM structures can indeed be modified by the addition or removal of functional groups, hydroxylation, methylation, glycosylation and acylation being the most frequent modification reactions.

Various SMs are widely accumulated in seeds of Brassicaceae model and crop species, which thus constitute an interesting model to study SMs diversity and functions (Corso et al., 2020; Corso et al., 2021; Baud et al., 2023). Major metabolic classes that are accumulated in seeds of Brassicaceae species, including the model species *Arabidopsis thaliana* and the crops *Camelina sativa* and *Brassica napus* (rapeseed), are the nitrogen-containing compounds glucosinolates, and many phenylpropanoids, including flavonols, proanthocyanidins (PAs, or condensed tannins), cinnamic acids and monolignols (Halkier and Gershenzon, 2006; Quéro et al., 2016; Barreda et al., 2024).

While some SM classes can be directly synthetized in seeds, others are synthetized in different organs or tissues of the mother plant and then transported to seeds. In *A. thaliana*, the flavonoids proanthocyanidins (PAs) and flavonols are synthetized in seeds (Lepiniec et al., 2006; Corso et al., 2020). In particular, the aglycone and decorated quercetin and kaempferol flavonols are accumulated at high concentration in the seed coat, endosperm and embryo. The localization of flavonols in *A. thaliana* seeds depends on their modifications, particularly on the glycosylation pattern, the monoglycosylated flavonols being accumulated in the seed coat and/or endosperm, and di and tri –glycosylated flavonoids accumulated in the embryo of Arabidopsis seeds (Routaboul et al., 2006). PAs are specifically synthetized in the seed coat, where they are accumulated at high concentrations at early Arabidopsis seed developmental stages, starting from 1-2 days after pollination (Lepiniec et al., 2006). Differently from flavonoids, glucosinolates (GSLs) are synthetized in maternal tissues outside of the seeds and then transported to seeds in *A. thaliana* (Nour-Eldin and Halkier, 2009; Sanden et al., 2023; Xu et al., 2023). Recent results nicely showed that, in *A. thaliana* seeds, GSL glucoiberin and glucoraphanin are synthetized in the funiculus and then transported to the seed embryo, where they accumulate and are subject to modifications (Sanden et al., 2023; Xu et al., 2023). In particular, the side chain of those two methionine-derived aliphatic GSL is hydroxylated by the 2-oxoglutarate-dependent dioxygenase AOP3 to form hydroxylated/hydroxylalkyl GSL (OH-GSL) (Kliebenstein et al., 2001). Previous reports suggested that GSL sinapoylation and benzoylation occurring on the -OH from the side chain structure are catalysed by SCPL17, a serine carboxypeptidase-like, while the synthesis of benzoyl compounds is catalysed by the benzoate-CoA ligase BZO1 (Reichelt et al., 2002; Kliebenstein et al., 2007; Lee et al., 2012; Sanden et al., 2023). Many abundant cinnamic acids in Brassicaceae seeds derive from sinapic acid core structure and decorated metabolites, mainly sinapate esters (COO-R) (Baumert et al., 2005; Corso et al., 2020). In *A. thaliana*, the synthesis of sinapate esters starts with the glycosylation of sinapic acid to form sinapoyl-glucose (sinapoyl-G), that constitutes the source of sinapate for the production of other sinapate-esters. These include the conversion of sinapoyl-G to sinapoyl-malate by the sinapoylglucose: malate sinapoyltransferase (SMT), the synthesis of 1,2-di-sinapoyl-G from two molecules of sinapoyl-G catalyzed by the sinapoyl-glucose:sinapoyl-glucose sinapoyltransferase or the synthesis of sinapoylcholine (sinapine) by 1-O-sinapoyl-beta-glucose:choline sinapoyltransferase (SCT) (Fraser et al., 2007; Stehle et al., 2008; Barreda et al., 2024).

While many of the above cited SMs are essential for Brassicaceae seeds to cope with abiotic and biotic stresses (Corso et al., 2020), many efforts have been done by breeders to drastically reduce their content in seeds since some of them are considered antinutritional compounds for human and animal consumption (Di Gioia et al., 2019). Among them sinapine (an alkaloidal amine) is one of the most abundant metabolites synthetized in Brassicaceae model and crop species and can constitute up to 2% of seed weight in these species (Menard et al., 2022). Sinapine reduces protein quality and digestibility and, because of its bitter taste, makes the meal obtained from Brassicaceae seeds less suitable and palatable for animal consumption (Bell, 1993; Huang et al., 2008; Menard et al., 2022). Similarly to other SMs accumulated in seeds, GSLs are part of the arsenal of compounds that plant species use to defend themselves against insects, vertebrate herbivores and pathogens (Dalling et al., 2020). GSLs, such as glucoraphanin, glucobrassicin and sinigrin, are considered antinutritional because of their bitter taste or their potential toxicity due to the accumulation of degradation products such as isothiocyanates, thiocyanates, nitriles and epithionitriles. Nevertheless, many reports highlighted a wide range of beneficial effects on human health, including but not exclusively due to their anticancerogenic, antibacterial, antidiabetic, and antioxidant activities (Di Gioia et al., 2019). In addition, despite GSLs were considerably reduced in Brassicaceae seed by breeding programs, limited information is available about the activity and/or toxicity of a large set of aglycone and decorated (e.g. glycosylated, sinapoylated and benzoylated) GSLs, together with their related degradation products.

Seeds from Brassicaceae crop species are used for oil and protein extraction, and for valorisation of co-products, including many SMs, but less information is available about SM diversity in these seeds and, hence, modifications, tissue-distribution and accumulation during seed development, unlike the extensive studies conducted in *A*. *thaliana*. Among these species *Camelina sativa* (camelina) is an oilseed crop cultivated for human and animal nutrition, and for industrial uses, increasingly used as model for research on Brassicaceae species. The characteristics of camelina make it a valuable crop for diversifying and intensifying production in agroecological cropping systems (Zubr, 1997; Neupane et al., 2022). Camelina seeds oil is rich in omega-3, polyunsaturated fatty acids (PUFA) and vitamin E (tocopherol), and is used in food, cosmetics and as a biofuel (Faure and Tepfer, 2016; Zanetti et al., 2017). Besides oil, camelina seeds accumulate high levels of SMs, including many phenylpropanoids and GSLs with a wide range of biological functions (Quéro et al., 2016; Alberghini et al., 2022; Boutet et al., 2022). *Camelina sativa* is hexaploid (2n=6x=40), with three sets of chromosomes and therefore three genomes (Kagale et al., 2014). *C. sativa* genome (n = 20, N^6^N^7^H) originated from a hybridization between the parental genomes *C. neglecta*–like (n = 13, N^6^N^7^) and *C. hispida* (n = 7, H) (Mandáková et al., 2020). In particular, every chromosome or chromosomal region of *A. thaliana*, was represented in three independent chromosomes in the genome of *C. sativa*. We recently analyzed the seed SM landscape in camelina seeds, highlighting the accumulation of a large set of SM classes, including GSLs, flavonoids, cinnamic acids, monolignols, terpenoids and amino acids and derivatives (Boutet et al., 2022). This work documented major effects of the environment on the stimulation of seed SMs in *C. sativa* seeds and showed that SMs, including GSLs and phenylpropanoids, showed a higher environmentally-induced plasticity with respect to most of the primary metabolites, including some sugars, fatty acids, proteins and lipids (Alberghini et al., 2022; Boutet et al., 2022). While we explored in detail SM diversity and plasticity, few information is available about SM distribution and expression of SM-related proteins and/or genes in developing and germinating seed tissues, including coat, endosperm and embryo, of camelina and other oilseed crop species (Fang et al., 2012). Knowing the spatiotemporal synthesis, transport and degradation of SMs in a crop species such as camelina would be indeed of main importance to develop new tools to finely modulate the ratio between beneficial and antinutritional compounds in seeds.

In this study we used untargeted metabolomics, proteomics and transcriptomics to analyze the synthesis, transport, modifications and degradations of SMs accumulated in the different seed tissues of *C. sativa* (i.e. seed coat, endosperm, and embryo), during seed development and germination. This multi-omic approach allowed to identify specific signatures of SMs, proteins and transcripts associated with the development of the camelina seed coat/endosperm and embryo. In addition, we highlighted contrasting seed composition and spatial distributions for the defence-related and antinutritional GSL compounds among closely related Brassicaceae species: Camelina, Arabidopsis and Brassica. While camelina plants produced long chain (C8-C11) GSL, Arabidopsis and Brassica accumulate mid- to short-chain (C3-C7) GSL (Czerniawski et al., 2021; Missinou et al., 2022). Our data further revealed specific GSL distributions within Brassicaceae seed tissues, with long chain methylsulfinylalkyl GSLs primarily accumulating in the seed coat and endosperm, while mid- to short-chain methylsulfinylalkyl GSLs were mainly localized in the embryo.

## Results

### *Camelina sativa* seed coat and embryo development

*Camelina sativa* seeds reach maturity in about 50 days from the onset of flowering. Plants were cultivated in a greenhouse (see materials and methods for more detailed information). Seeds were harvested at 13, 21, 28, 35, 42 Days After Flowering (DAF), at dry seeds, and at two germination stages (6 hours and 24 hours from seed imbibition) (Figure 1). Seed coat and endosperm (SCE), and seed embryo (SE) were separated and used for multi-omic analyses. As for the ratio between the two seed tissues, at 13 DAF 77% of the seed weight is constituted by the SCE, at 21 DAF the two tissues represent together 50% of the seed weight, while from 28 DAF the SE represents around 70% of the seed weight (Figure 1).

**Figure 1.**
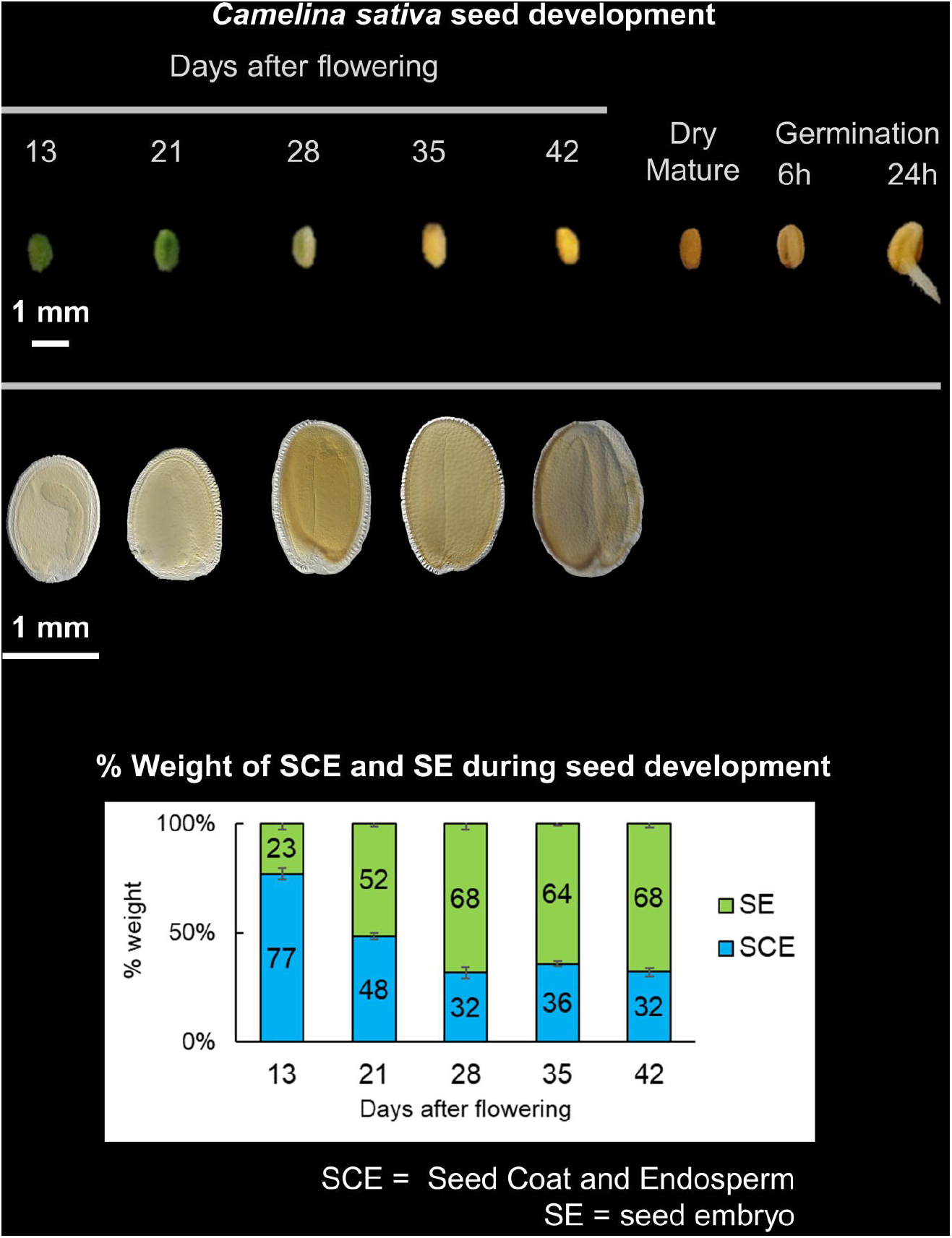
*Camelina sativa* seeds used in this study. Seed coat and embryo samples used in this study were harvested at 13, 21, 28, 35, 42 days after flowering. Dry mature and germinating seeds (6h and 24h) were also used.

### Specialized metabolite diversity in camelina seed coats and embryos

Untargeted SM profiling revealed a large diversity in camelina seeds (Figure 2; Table S1 and S2A). Untargeted metabolomic analyses (positive and negative ElectroSpray Ionization, ESI+ and ESI-) allowed the detection of 4061 peaks of putative metabolites (i.e. metabolic features) (Table S2B). Molecular network analyses enabled the clustering of SM compounds according to their MS/MS spectra (Figure 2A, Table S2A) and was used to improve the annotation of the unknown metabolic features (Olivon et al., 2018; Boutet et al., 2022). Annotated SMs in camelina seed coats and embryos belong to several classes, among which amino acids (AA) and derivatives (208 SMs), cinnamic acids (176 SMs) and flavonols (64 SMs) were the most numerous, followed by nucleosides and derivatives, terpenoids, flavan-3-ols and proanthocyanidins (PAs), fatty amides, carbohydrates, carboxylic acids, glucosinolates (GSLs), flavonoids, sulfoxides, lignans, glutathione and derivatives, isothiocyanates (ITCs) and amines (Figure 2B and Table S2A).

**Figure 2.**
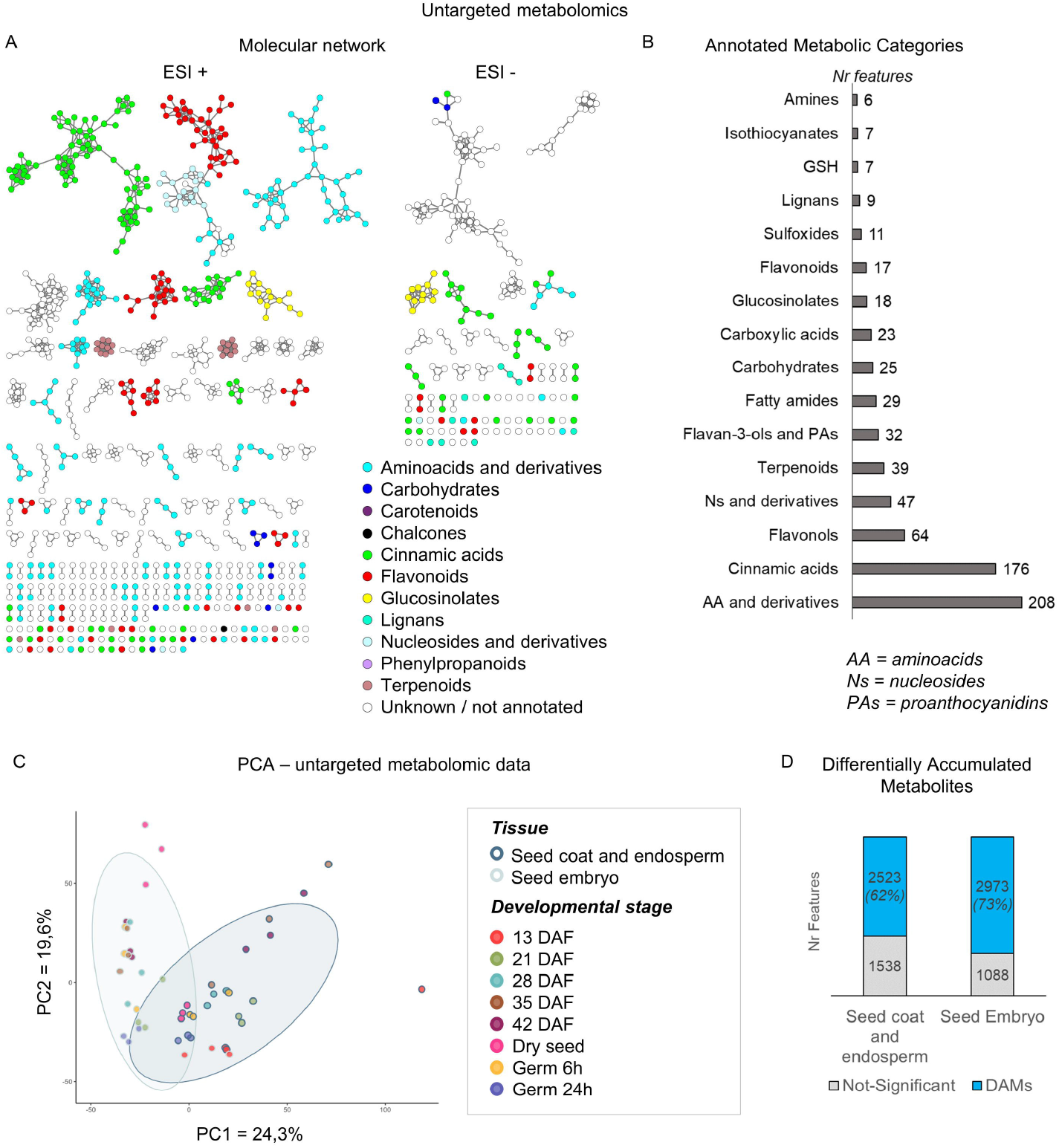
Untargeted metabolomics in developing and germinating SE and SCE. **A)** The molecular networks are shown for metabolite analyses carried out in positive (ESI+) and negative (ESI-) ionization modes. Different colours correspond to different metabolic classes. Metabolites are grouped based on their chemical structures. Cosine similarity scores of 0.85 and 0.65 were used for ESI+ and ESI-, respectively. **B)** Histogram showing the annotated/known metabolic categories. The number of metabolites that belong to each category is indicated in the figure. Categories with less than two metabolite features are not shown. **C)** Principal component analyses using the untargeted metabolomic data of SCE and SE harvested at several developmental stages and during germination. DAF = Days After Flowering. **D)** Venn diagram indicating the number of Differentially Accumulated Metabolite features (DAMf) according to the tissue and developmental stage factors.

The average intensity of SMs in seed coats and embryos allowed the identification of the top-10 metabolites with highest relative intensity (Table S1). Sinapine (cinnamic acids/alkaloids) was the SM showing the highest intensity for both seed coat/endosperm (SCE) and embryos (SE). In addition, two ITCs (9-Methylsulfinylnonyl ITC and 10-Methylsulfinyldecyl ITC) deriving from the degradation of two major GSL compounds, rutin (flavonol) and adenosine (nucleosides and derivatives) were highly accumulated metabolites common to seed coats and embryos. Among the SMs with high intensity and specific to the SCE, there were one ITC (11-Methylsulfinylundecyl ITC), the GSL glucoarabin (9-methylsulfinylnonyl GSL) and a GSL derivative (p2553; proposed formula: C_17_H_32_NO_7_S_2_^+^), together with the epicatechin (flavan-3-ol) and a cinnamic acid with unknown formula (p8). Among the seed-embryo–specific SMs with the highest intensity there were a triglycosylated flavonol, putatively annotated as quercetin-(xyloside-rhamnoside-glucoside), a sulfoxide (p29, proposed formula C_13_H_25_NOS) and three AA and derivatives (gamma-Glu-Leu, p13 with proposed formula C_11_H_20_N_2_O_5_S and (2S)-2-amino-5-[[(1S)-1-carboxy-2-methylsulfanylethyl]amino]-5-oxopentanoic acid).

### Camelina seed specialized metabolome is shaped by the tissue and the developmental stage

The untargeted metabolomic data obtained from seed coats and embryos harvested at six developmental and two germination stages were used to perform a principal component analysis (PCA). 24,3 % and 19,6 % of the variability observed among the samples can be explained respectively by the axis PC1, which is associated to the type of tissue, and the axis PC2, which is associated to the developmental stage (Figure 2C). The PCA performed allowed to clearly separate the samples according to the tissue, confirming that seed coat/endosperm and embryo present contrasting specialized metabolome signatures. Hence, Differentially Accumulated Metabolites (DAMs) according to the “developmental stage” (13 days after flowering (DAF) to dry seeds, and germinating seeds) factor were identified (ANOVA, FDR<0.05) separately for the two seed tissues considered (SCE and SE). The statistical analysis highlighted that the accumulation of 62% (2523) and 73% (2973) of SMs was impacted by the seed developmental stage in the SCE and SE, respectively (Figure 2D).

### A core-set of specialized metabolites is regulated during seed embryo and coat-endosperm development and maturation

In order to identify stage-specific DAMs, a hierarchical clustering analysis was conducted for SCE and SE tissues using DAMs intensities. The analysis allowed the separation of the DAMs into five clusters for seed coat-endosperm (I to V; Figure 3A) and four clusters for seed embryo (VI to IX; Figure 3B). The clusters allowed the identification of DAMs accumulated from early (13 DAF) to late (42 DAF and dry seeds) seed developmental stages, and of germination-specific DAMs. Afterwards, using the DAMs belonging to each cluster, an enrichment analysis was performed to identify the metabolic categories that are characteristics of each tissue and developmental stage. The enrichment analysis was performed using the metabolic categories identified with the molecular network analysis, both the ‘annotated’ and ‘not-annotated’ (or ‘unknown’) ones (Table S2A, column “Metabolite class”). In particular, not-annotated metabolites that clustered together in the molecular network analysis (Figure 2A) and therefore have similar chemical structures, were regarded as a distinct metabolic category. Besides the SMs assigned to known metabolic categories, several not-annotated SMs showing similar MS/MS profiles were enriched in different clusters displaying specificities to some seed tissue and/or developmental stage (Figure 3A and B).

**Figure 3.**
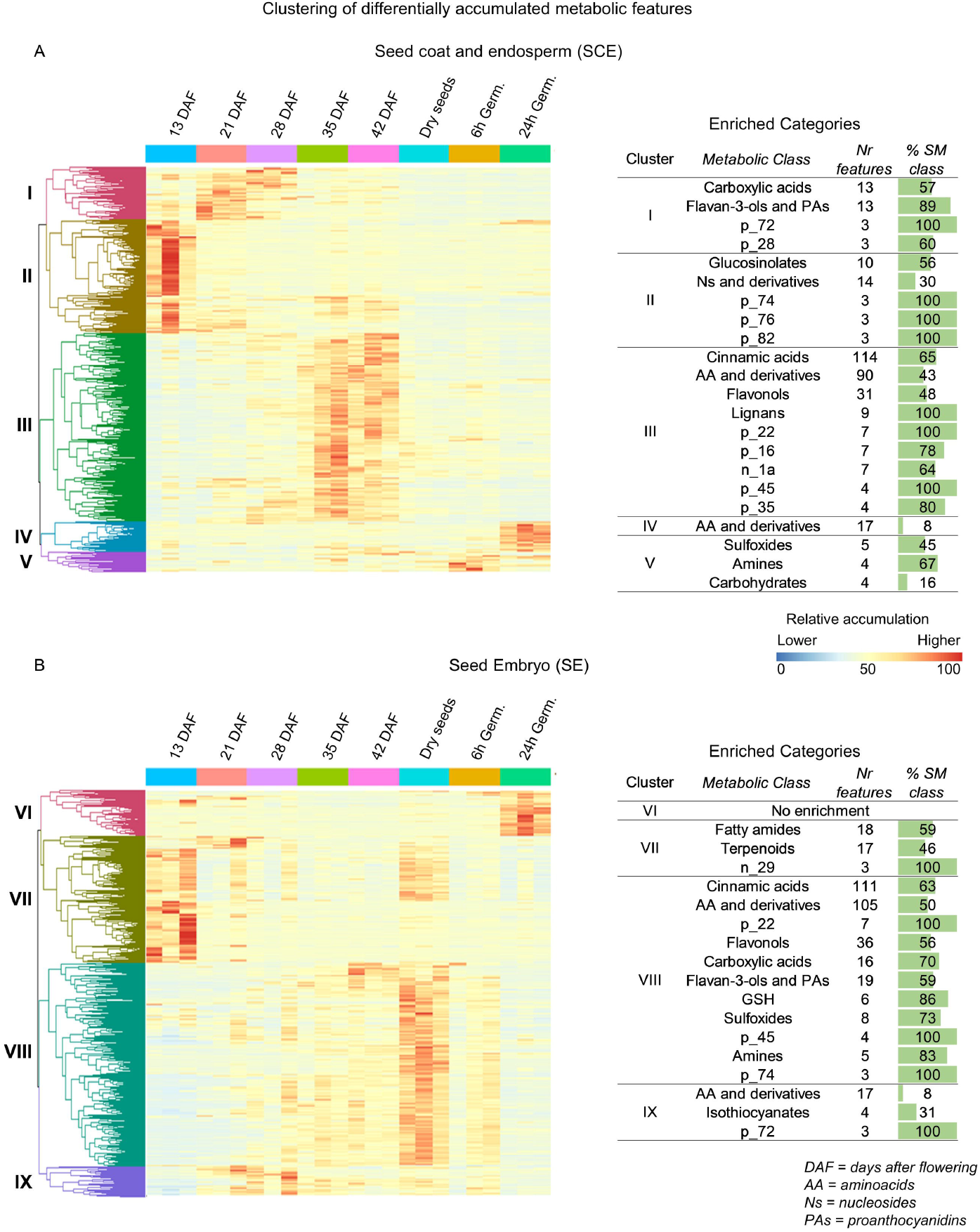
Specialized metabolites signatures and stage-specific enrichment of metabolic categories during SE and SCE development and germination. Hierarchical clustering and enrichment analyses of Differentially Accumulated Metabolites (DAMs) specific to SCE **(A)** and SE **(B).** Annotated and unknown enriched (hypergeometric test, p < 0.05) metabolic categories of DAMf clusters specific to seed coats and embryos are showed.

Regarding the SCE specialized metabolome (Figure 3A), carboxylic acids and flavan-3-ols and PAs were found to be enriched and specific to cluster I, characterized by DAMs with higher accumulations at 21 DAF and 28 DAF. In addition, glucosinolates, and nucleosides and derivatives SMs were found to be enriched in cluster II, which included DAMs with high intensity at 13 DAF. Cinnamic acids, AA and derivatives, flavonols and lignans categories were enriched in cluster III, which was the biggest for SCE tissues and showed SMs with increasing accumulation from 28 to 42 DAF. Finally, AA and derivatives, sulfoxides, amines and carbohydrates were enriched in clusters IV and V that are specific to seeds during germination stages.

Concerning the seed embryo (SE) SMs (Figure 3B), no enriched metabolic categories were found for SMs from cluster VI, which is characterized by DAMs more accumulated during germination 24h after imbibition. Cluster VII, which includes DAMs with the highest accumulation mainly at the early seed developmental stage (13 DAF), is enriched for fatty amides and terpenoids. Cluster VIII is the largest one and contains SMs with increasing intensity from 35 DAF to dry seeds, and showed an enrichment for several metabolic classes, including cinnamic acids, AA and derivatives, flavonols, carboxylic acids and flavan-3-ols and PAs. From work on Arabidopsis we know that flavan-3-ols and derivatives are not produced in the embryo (Lepiniec et al., 2006) and that these metabolites could originate from degraded cells of the endothelium and, hence, will not be further discussed in this article. Finally, AA and derivatives and isothiocyanates (GSLs degradation products) were found to be enriched in cluster IX, which is characterized by DAMs with higher accumulation predominantly at 21 and 28 DAF. According to the results highlighted by the enrichment analysis and the broad interest for seed biology and quality, we will focus on phenylpropanoid (flavonoids, cinnamic acids and monolignols) and GSL metabolic categories, and on genes or enzyme categories involved in their modifications (i.e. glycosylation, methylation, hydroxylation and acylation) (Barreda et al., 2024).

### Proteomic analyses allowed the identification of SM-related proteins with specific expression pattern during seed development

To get a better understanding of the regulation and accumulation dynamics of SM-related pathways in camelina SCE and SE, data-independent acquisition (DIA) proteomic combined with ion mobility spectrometry analyses were performed on the same samples used for untargeted metabolomics. These analyses allowed to measure the abundance of 11462 proteins (Table S3). The PCA performed with proteomic data separated the samples according to seed developmental and germination stages (Figure 4A). As for metabolomic data, Differentially Accumulated Proteins (DAPs) impacted by the “developmental stage” factor were identified (ANOVA, FDR<0.05) separately for SCE and SE (Figure S1). The statistical analysis highlighted that the accumulation of 14% (995) and 33% (2418) DAPs was specifically affected by the seed developmental stage in SCE and SE, respectively, while 53% (3912) of detected proteins were differentially expressed and common to SCE and SE (Figure 4B and S1). Next, the proteins involved in GSLs, flavonoids, cinnamic acids and monolignols biosynthesis/pathways, and the enzymes putatively involved in SM modifications that were differentially expressed in SCE (333 DAPs) and SE (374 DAPs) during development, were identified. Among these DAPs, 273 were in common between SCE and SE (Table S3 and Figure 5). A hierarchical clustering analysis was conducted for SCE and SE seed tissues using GSLs, flavonoids, cinnamic acids and monolignols –related DAPs (Figure 5A and 5B). DAPs were separated into six clusters for SCE (Figure 5A) and seven clusters for seed embryo (Figure 5B), according to their accumulation patterns. In SCE, most DAPs belonging to selected SM categories were accumulated at earlier seed developmental stages, either at 13 and/or 21 DAF (clusters IV and VI). Similar results were observed for SE, except for proteins involved in SM glycosylation (belonging to glycosyltransferase GT1 group) that were accumulated at several seed developmental and maturation stages, particularly during seed germination (cluster VII).

**Figure 4.**
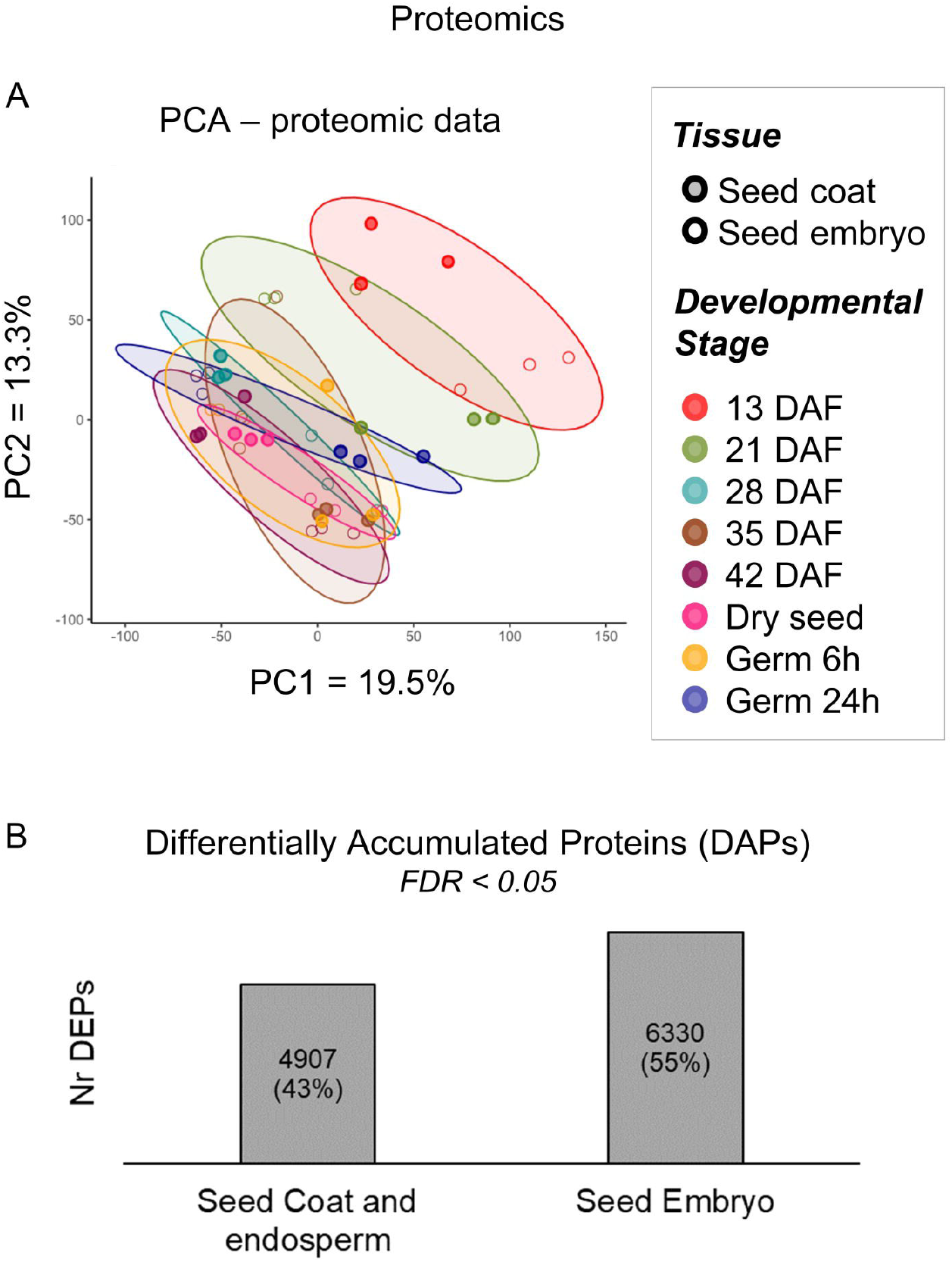
Proteomic analyses in developing SCE and SE. **A)** Principal component analyses using the proteomic data of SCE and SE harvested at several developmental stages and during germination. DAF = Days After Flowering. **B)** Histogram indicating the number of Differentially accumulated proteins (DAPs) according to the developmental stage in seed coat and embryo.

**Figure 5.**
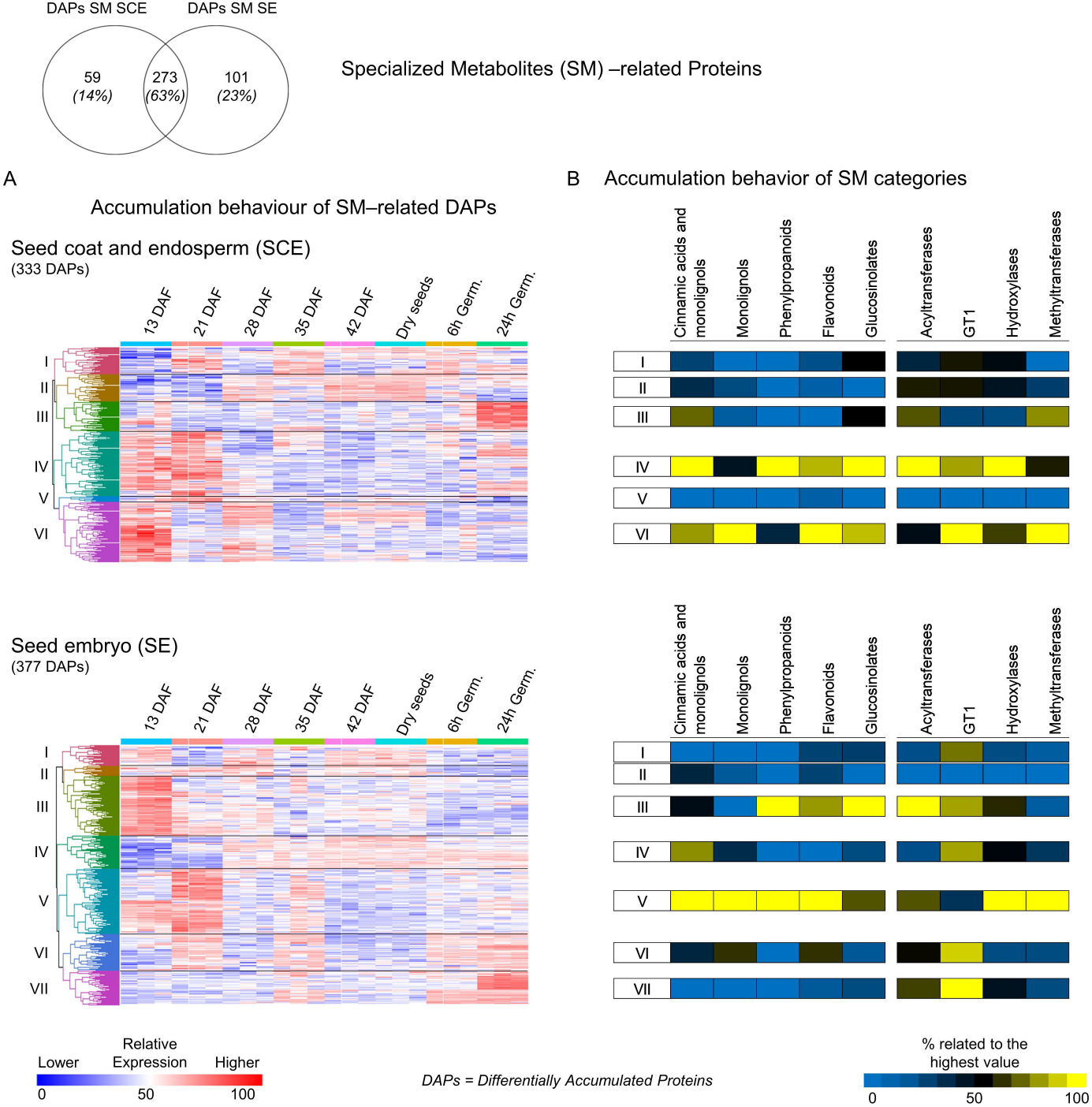
Accumulation pattern of specialized metabolites-related proteins. A) Hierarchical clustering of SM-related DAPs in SCE and SE. **B)** Expression behaviour of proteins involved in the biosynthesis, transport and degradation of the major specialized metabolite pathways, and of the main SM-decoration/modification categories. GT1 = glycosyltransferases, group 1.

### Multi-omic data revealed contrasting glucosinolate landscapes and spatial distribution in seeds of camelina and other Brassicaceae species

Glucosinolates and their related degradation products were further characterized in Camelina SE and SCE (Figures 6, 7). The results revealed that GSLs and their degradation products are differentially accumulated in the SCE and/or the SE of camelina seeds (Figure 7). Most GSLs that were differentially accumulated during seed development, including the seed-specific aliphatic GSLs 8-methyl-sulfinyl-octyl-GSL, glucoarabin (9-Methylsulfinylnonyl-GSL; GSL9), glucocamelinin (10-Methylsulfinyldecyl-GSL; GSL10) and gluconesliapaniculatin (11-Methylsulfinylundecyl-GSL; GSL11), were strongly accumulated in the SCE and showed the highest accumulation level at early seed developmental stage (13 DAF) and/or at later stage (35 DAF) (Figure 7A). The indolic GSLs glucobrassicin and neoglucobrassicin showed their highest accumulation at 13 DAF. In contrast, most of the GLS degradation products (ITCs) were differentially accumulated only in camelina embryos, where they reached their peak levels at dry mature seed stage in the SE (Figure 7A, 7B). The contrasting GSL distributions between the SCE and SE is further highlighted by comparing the relative accumulation levels of the main annotated and differentially accumulated GSLs and their ITC degradation products in these tissues (Figure 7B). Glucoarabin (GSL9) was the GSL with the highest relative abundance, peaking at 35 DAF in SCE. Consistently, the main GSL degradation product, 9-Methylsulfinylnonyl ITC, reached its maximum relative intensity at 13 DAF in the SCE and in the dry seed embryo. Gluconesliapaniculatin (GSL11) was the second most abundant GSL, with its peaked at 35 DAF, followed by glucocamelinin (GSL10), which was characterized by two peaks at 13 and 35 DAF (Figure 7B). The corresponding degradation products of GSL10 and GSL11,10-Methylsulfinyldecyl and 11-Methylsulfinylundecyl ITCs, were differentially and highly accumulated in the SE of dry mature seeds (Figure 7B). Finally, 8-Methylsulfinyloctylglucosinolate was also specific to the SCE, but showed lower intensity compared to GSL9, GSL10 and GSL11.

**Figure 6.**
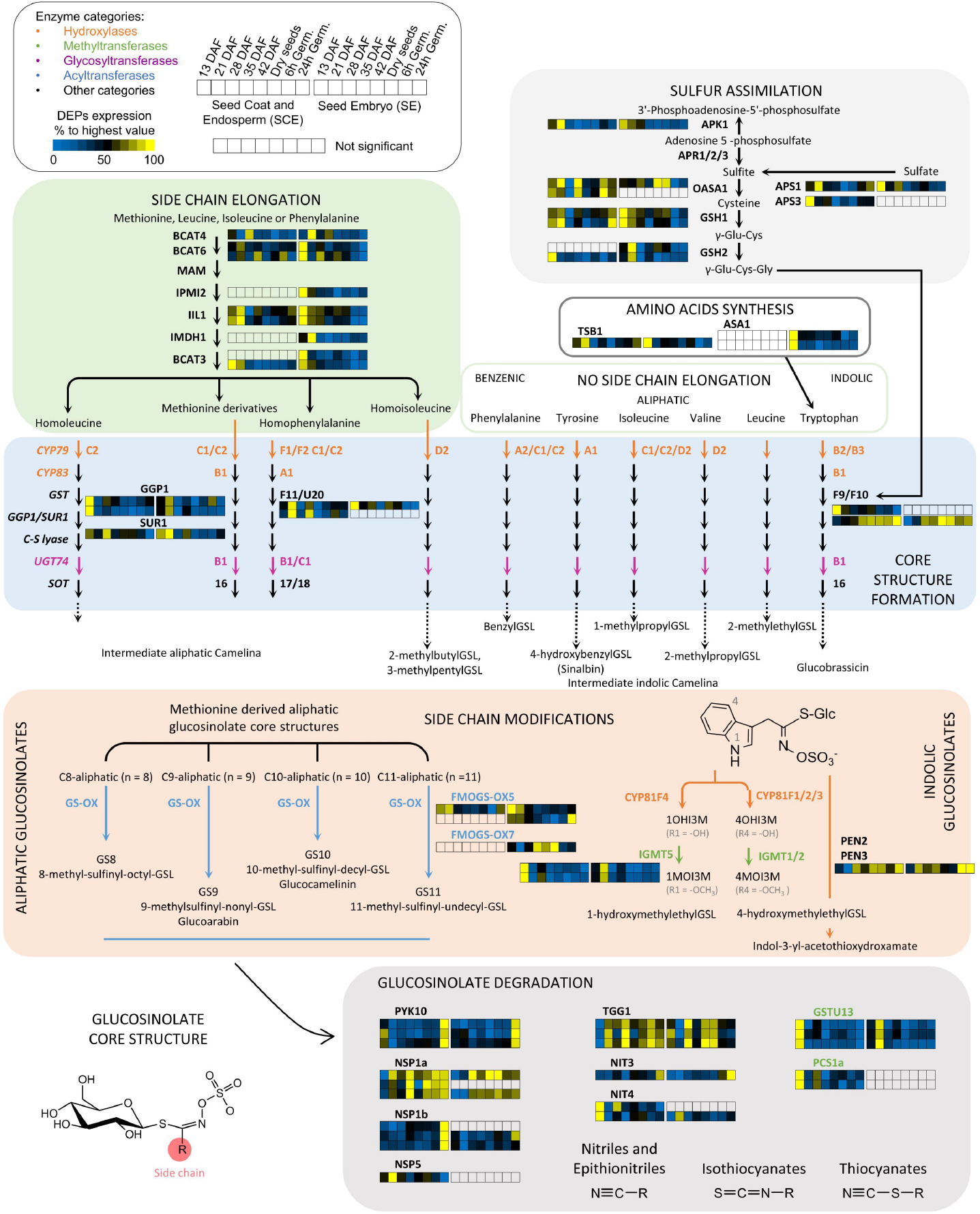
Expression of proteins involved in glucosinolate pathways. Expression of proteins involved in GSL biosynthesis, side-chain elongation and modifications, and degradation in camelina SCE and SE. Only differentially accumulated proteins (DAPs, ANOVA, FDR < 0.05) are showed in the figure.

**Figure 7.**
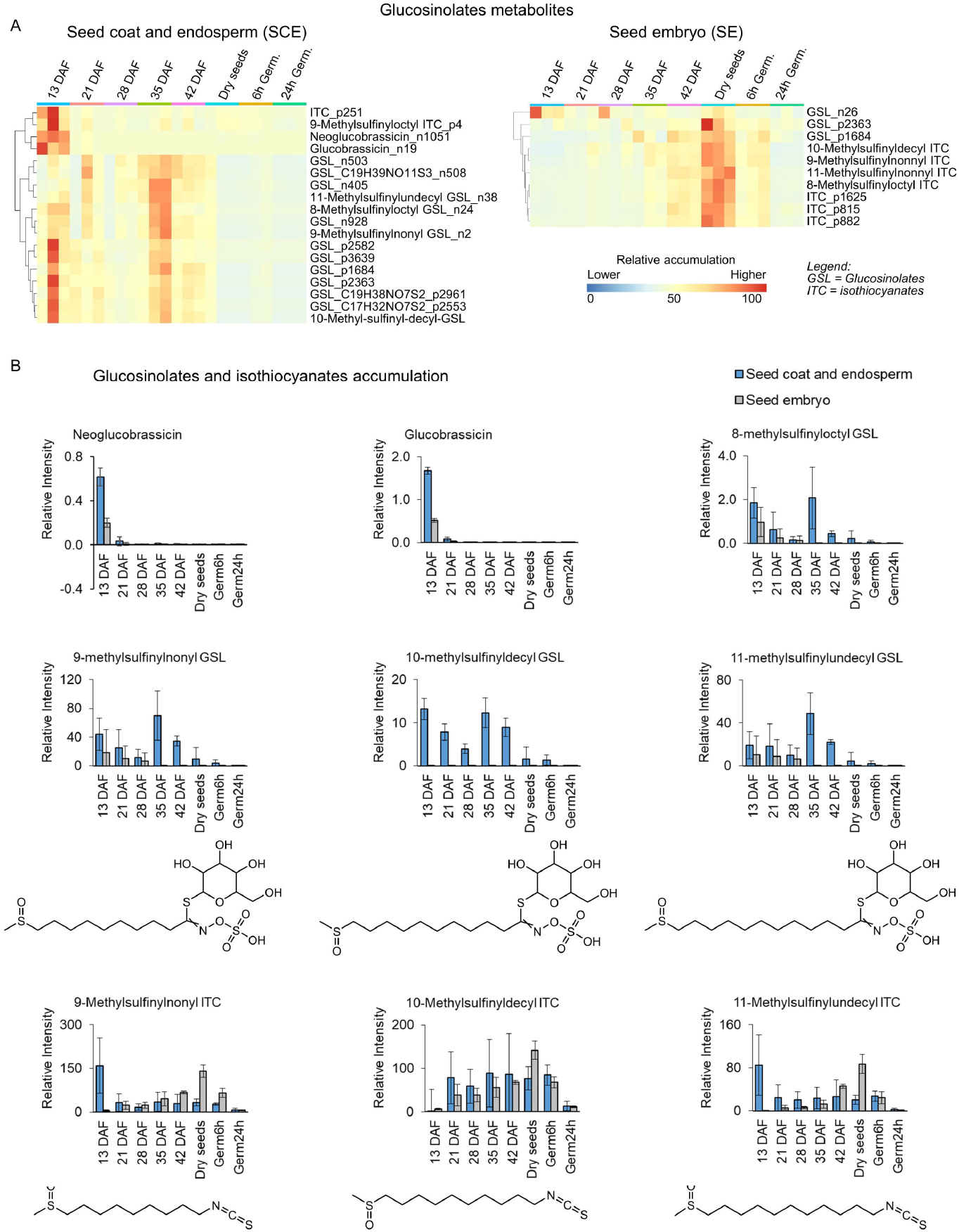
Accumulation profiles of glucosinolates and isothiocyanates metabolites in SCE and SE. **A)** Heatmap showing the glucosinolates and isothiocyanates degradation products that are differentially accumulated during seed development in SCE and SE tissues. **B)** Histograms showing the major glucosinolates and isothiocyanates specialized metabolites accumulated in SCE and/or SE.

In most cases, the accumulation of proteins involved in GSL side-chain elongation and core-structure formation was higher at earlier seed developmental stages (13 and 21 DAF) in both SCE and SE (Figure 6). Among the proteins related to GSL modifications, we identified two Flavin-Monooxygenases, namely FMOGS-OX5 and FMOGS-OX7, putatively implicated in GSL hydroxylation. These proteins showed the highest accumulation at early (FMOGS-OX5, 13-21 DAF) or late (FMOGS-OX7, 35 DAF to dry seeds, specific to SE) seed developmental stages. FMOGS-OX5 was also found in the SE during germination. Proteins playing a role in GSL degradation were accumulated during the whole seed development and germination in both SCE and SE, with some specificities and exceptions. For example, PYK10 myrosinase showed the highest accumulation during germination, while TGG1 myrosinase was accumulated throughout seed development in both SCE and SE, and also during germination specifically in SCE tissues (Figure 6).

These results mainly highlighted that in *C. sativa* GSLs compounds were unexpectedly accumulated in both the seed coat and endosperm, while their corresponding degradation products, ITCs, were present at high level in the embryo. Next, GSL and ITC accumulations were characterized in the seed coats/endosperms and embryos of two others Brassicaceae species, *A. thaliana* and *B. napus*. These results showed that camelina seeds are characterized by different GSL accumulation patterns and expression of related genes/proteins compared to *A. thaliana* and *B. napus* (Figure 8a; Table S4). **First**, we showed that *C. sativa* exclusively accumulated long carbon(C)-chain GSLs, notably C9-C11 methylsulfinylalkyl GSLs, in the seed coat and the endosperm, while Arabidopsis and Brassica seeds did not synthetize these compounds (Matsuda et al., 2010; Missinou et al., 2022). **Second**, we showed that the long chain C8 methylsulfinylalkyl GSLs are accumulated in the seed coat and endosperm of *C. sativa* and in *A. thaliana*, but are not synthetized and accumulated in *B. napus* seeds. **Third**, mid- and short-chain C3-C7 methylsulfinylalkyl GSLs are accumulated in the embryo of *B. napus* and in *A. thaliana*, but are not synthetized and accumulated in *C. sativa* seeds. **Fourth**, ITCs (GSL degradation products) are accumulated in the embryo in *C. sativa* and in *B. napus*, and in the seed coat and endosperm in *A. thaliana*. **Fifth**, benzoylated and sinapoylated GSLs were found exclusively in *A. thaliana* seeds. In accordance with this finding, no Camelina orthologue of the SCPL17 acyltransferase enzyme, involved in GSL side-chain benzoylation and sinapoylation in *A. thaliana*, has been found in our proteomic data (Figure 6, Table S3 and Table S5). These multi-omic data revealed a complex and fine-tuned regulation of GSL synthesis, modification and transport that has been summarized in Figure 8b.

**Figure 8.**
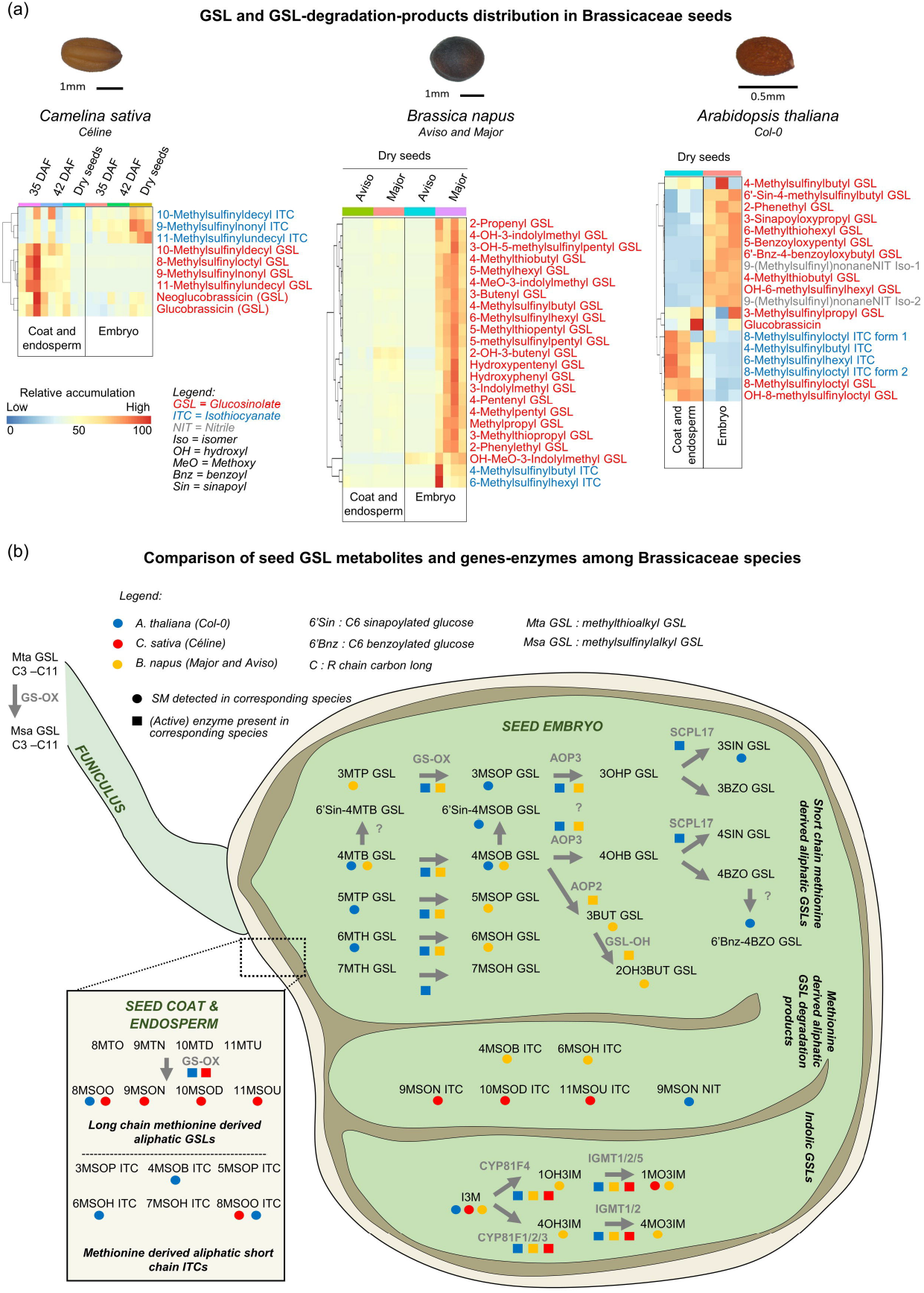
Distribution of glucosinolates and related degradation products in different Brassicaceae species. **A)** Heatmap showing the glucosinolates and related degradation products present in *Camelina sativa*, *Brassica napus* and *Arabidopsis thaliana* seed coat/endosperm and embryo tissues. **B)** Model summarizing GSL metabolites distribution and expression of related enzymes in seed tissues of *C. sativa*, *B. napus* and *A. thaliana*. Abbreviations: I3M, 3-indolylmethyl GSL (Glucobrassicin); 1OHI3M GSL, 1-hydroxyindol-3-ylmethyl GSL; 4OHI3M GSL, 4-hydroxyindol-3-ylmethyl GSL; 1MOI3M GSL, 1-methoxyindole-3-ylmethyl GSL (Neoglucobrassicin); 4MOI3M GSL, 4-methoxyindole-3-ylmethyl GSL; 3MTP GSL, 3-methylthiopropyl GSL (Glucoiberverin); 4MTB GSL, 4-methylthiobutyl GSL (Glucoerucin); 5MTP GSL, 5-methylthiopentyl GSL (Glucoberteroin); 6MTH GSL, 6-methylthiohexyl GSL (Glucolesquerellin); 7MTH GSL, 7-methylthioheptyl GSL; 8MTO GSL, 8-methylthiooctyl GSL; 9MTN GSL, 9-methylthiononyl GSL; 10MTD GSL, 10-methylthioecyl GSL; 11MTU GSL, 11-methylthioundecyl GSL; 6’Sin-4MTB, 6’sinapoyl-4-methylthiobutyl GSL; 3MSOP GSL, 3-methylsulfinylpropyl GSL (Glucoiberin); 4MSOB GSL, 4-methylsulfinylbutyl GSL (Glucoraphanin); 5MSOP GSL, 5-methylsulfinylpentyl GSL (Glucoalyssin); 6MSOH GSL, 6-methylsulfinylhexyl GSL (Glucohesperin); 7MSOH GSL, 7-methylsulfinylheptyl GSL (Glucoibarin); 8MSOO GSL, 8-methylsulfinyloctyl GSL (Glucohirsutin); 9MSON GSL, 9-methylsulfinylnonyl GSL; 10MSOD GSL, 10-methylsulfinyldecyl GSL; 11MSOU GSL, 11-methylsulfinylundecyl GSL; 6’Sin-4MSOB, 6’sinapoyl-4-methylsulfinyl GSL; 3OHP GSL, 3-hydroxypropyl GSL; 4OHB GSL, 4-hydroxybutyl GSL; 3BUT GSL, 3-butenyl GSL (Gluconapin); 2OH3BUT GSL, (2R)-2-hydroxy 3-butenyl GSL (Progoitrin); 3BZO GSL, 3-benzoyloxypropyl GSL (Glucomalcomiin); 3SIN GSL, 3-sinapoyloxypropyl GSL; 4BZO GSL, 4-benzoyloxybutyl GSL; 4SIN GSL, 4-sinapoyloxybutyl GSL; 6’Bnz-4BZO GSL, 6’benzoyl-4-benzoyloxybutyl GSL; 3MSOP ITC, 3-methylsulfinylpropyl ITC (Iberin); 4MSOB ITC, 4-methylsulfinylbutyl ITC (Sulforaphane); 5MSOP ITC, 5-methylsulfinylpentyl ITC; 6MSOH ITC; 6-methylsulfinylhexyl ITC; 7MSOH ITC, 7-methylsulfinylheptyl ITC; 8MSOO ITC; 8-methylsulfinyloctyl ITC; 9MSON ITC; 9-methylsulfinylnonyl ITC; 10MSOD ITC, 10-methylsulfinyldecyl ITC; 11MSOU ITC, 11-methylsulfinylundecyl ITC; 9MSON NIT, 9-methylsulfinylnonylnitrile.

### Multi-omic data highlighted seed coat-endosperm and embryo-specific accumulation patterns of proteins and metabolites related to glucosinolates, flavonoids and cinnamic acids

To get a better overview about other major SM categories, the accumulations of metabolites and proteins belonging to flavonoids (Figures 9) and cinnamic acids and monolignols (Figures S2 and S3) categories were further analysed in camelina seed tissues. Given that *C. sativa* has a hexaploid genome, three homeolog genes could potentially encode proteins involved in a specific enzymatic activity. The proteomic results showed in this study allowed the identification of the proteins copies specifically accumulated in SE and SCE and involved in SM biosynthesis, modifications and degradation (Table S3; Figure 5). For GSLs, cinnamic acids and monolignols, one gene copy was expressed for most proteins involved in the same enzymatic activity in both SCE and SE. In contrast, flavonoid-related proteins were mostly accumulated with one gene copy for SCE and two copies for SE (Figure S4).

**Figure 9.**
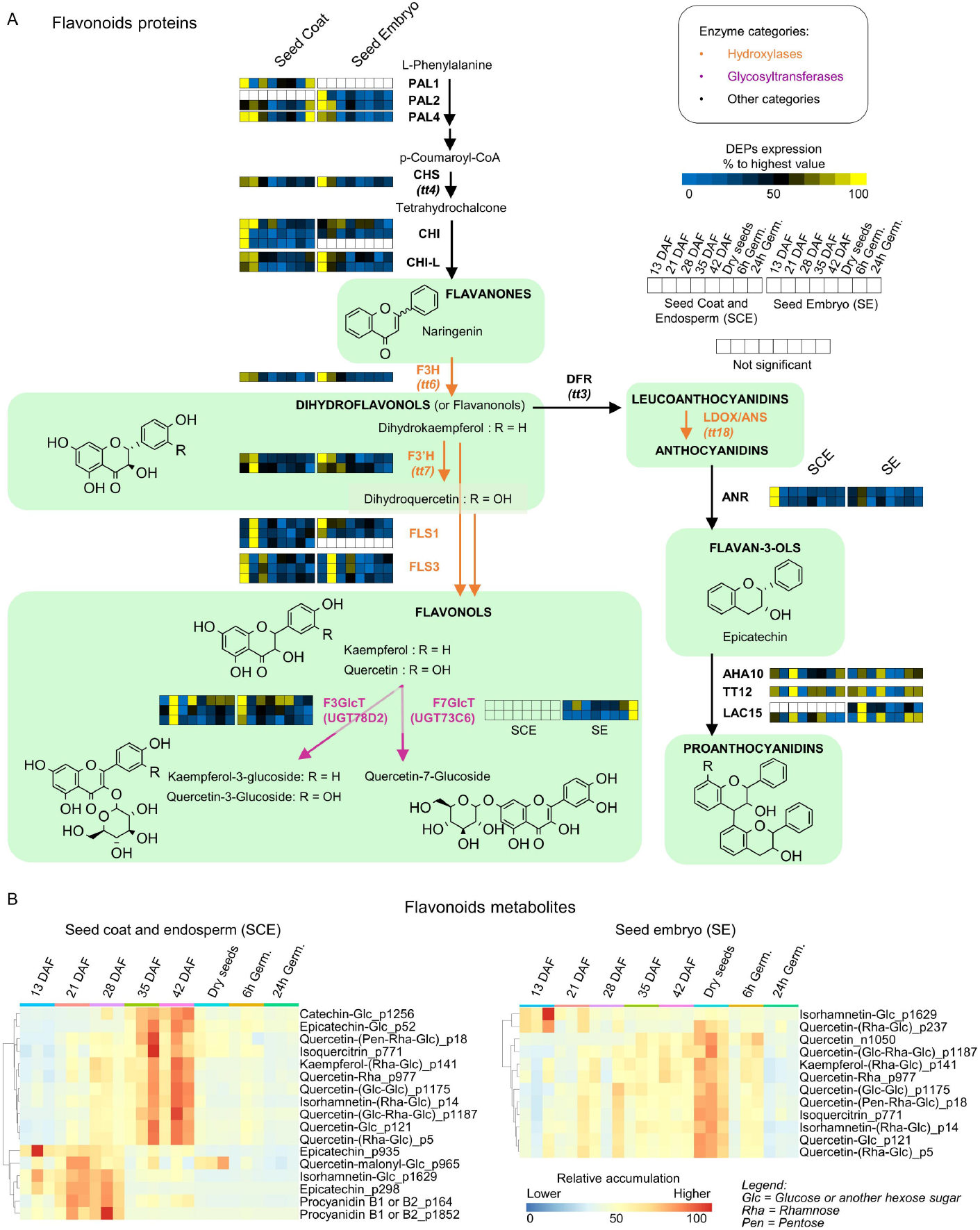
Accumulation profiles of flavonoids proteins and metabolites in SCE and SE. **A)** Expression of proteins involved in flavonoid biosynthesis, degradation, transport and glycosylation in camelina SCE and SE. Only differentially accumulated proteins (DAPs, ANOVA, FDR < 0.05) are showed in the figure. **B)** Heatmap showing flavonoid metabolites that are differentially accumulated during seed development in SCE and SE tissues.

As observed for GSLs, the accumulation of different flavonoid classes showed marked differences depending on the seed tissue and the developmental stage (Figure 9B). In particular, in SCE tissue flavan-3-ols and derivatives were accumulated from 13 to 28 DAF in their aglycone form (epicatechin, catechin and procyanidin B1 and B2), while glycosylated flavan-3-ols (epicatechin and catechin glycosides) showed the highest accumulation level at 35 and 42 DAF. Most flavonol metabolites, including quercetin and kaempferol mono-, di- and tri-glycosides, were accumulated at later seed developmental stages in SCE (35-42 DAF) and SE (dry mature seeds). In contrast, isorhamnetin (i.e. methylated quercetin)-glycoside was predominantly accumulated at earlier developmental stages in both tissues (Figure 9B). Interestingly, when the isorhamnetin presented a second-glycosyl group (isorhamnetin-(Rha-Glc), it was mainly accumulated at later seed developmental stages in both tissues (Figure 9B). Enzymes involved in the biosynthesis of the core flavonoid pathway and in flavonol specific pathway were expressed at 13-21 DAF in both SCE and SE, with a shift between the two tissues for some of them (Figure 7). This is the case of flavonol synthase 1 (FLS1) which showed the highest expression at 21 DAF in the SCE and at 13 DAF in the SE. On the contrary, flavonol synthase 3 (FLS3), showed an earlier expression in SCE compared to the SE. Anthocyanidin reductase (ANR), involved in flavan-3-ols synthesis showed its highest expression in the SCE at early developmental stages. As concerns proteins involved in flavonoid glycosylation, it is worth mentioning that the UDP-dependent glycosyltransferase (UGT), F7GlcT, was specific to the embryo and showed the highest expression during the germination stage.

While the metabolite landscape and composition of GSLs and flavonoids differed markedly between the SCE and SE, fewer differences between in cinnamic acid composition were observed between these two tissues (Figure S3). Sinapine was the most abundant metabolite belonging to this category in both tissues (Table S1), and showed the highest intensity at 35-42 DAF for SCE and in dry seeds for SE. Similarly, other cinnamic acid derivatives, e.g. sinapoyl, feruloyl and coumaroyl glycosides, were accumulated at 35-42 DAF in the SCE and at dry seeds in the embryo. In addition, guaiacyl-ferulic acid-glucose was more abundant during germination in SCE and SE. Some differentially accumulated proteins involved in cinnamic acids synthesis or decoration showed specific expression in seed embryo (Figure S2). This is the case of the sinapic acid:UDP-glucose glucosyltransferase (SGT1), involved in sinapic acid glycosylation, of the Sinapoylglucose:malate sinapoyltransferase (SMT), which produced sinapoyl-malate, and of 1-O-sinapoyl-beta-glucose:choline sinapoyltransferase, catalysing the synthesis of sinapine.

### Transcriptomic analyses in seed embryo

Transcriptomics were performed on a subset of samples compared to those used for metabolomic and proteomic analyses. In particular, seed embryo at 13, 21, 28, 35, 42 DAF and at 6 hours after germination stage were subjected to RNA-Seq analyses (Figure 10, Figure S5 and Table S5). As observed for proteomic and untargeted metabolomic analyses, large differences in gene expression (FDR < 0.05) were observed between seed developmental stages, and between seed development at 42 DAF and at 6 hours of germination. The PCA confirms these results and the quality of the data obtained since it allowed a clear separation of embryos from different developmental stages and showed that the three biological replicates were clustered together (Figure 10). Next, we extracted Differentially Expressed Genes (DEGs) involved in GSL, flavonoid, and cinnamic acid synthesis, transport, decorations (or modifications), degradation and regulation (Figure 10 and Figure S6). Many DEGs related to GSLs were expressed at 13 DAF and showed generally an earlier expression peak compared to what has been observed for corresponding proteomic data. As for flavonoids, all DEGs involved in their biosynthesis and most DEGs involved in their decoration showed the highest expression level at 13 DAF. *MYB11* and *MYB111* transcription factors (TFs) involved in the regulation of flavonol metabolites, were expressed at 13DAF, while *TTG1* TF showed its expression peak at germination stage. DEGs related to cinnamic acids were characterized by larger differences concerning their expression pattern. In particular, genes coding for Serine-carboxypeptidase-like enzymes involved in sinapic acid ester biosynthesis showed the highest expression level at 13 DAF (*SCPL12-FTP1*), 21 DAF (*SCPL9-SNG2-SCT*), 28 DAF (*SCPL7*), 42 DAF (*SGT1*) and during germination for *SST* (*SCPL9*).

**Figure 10.**
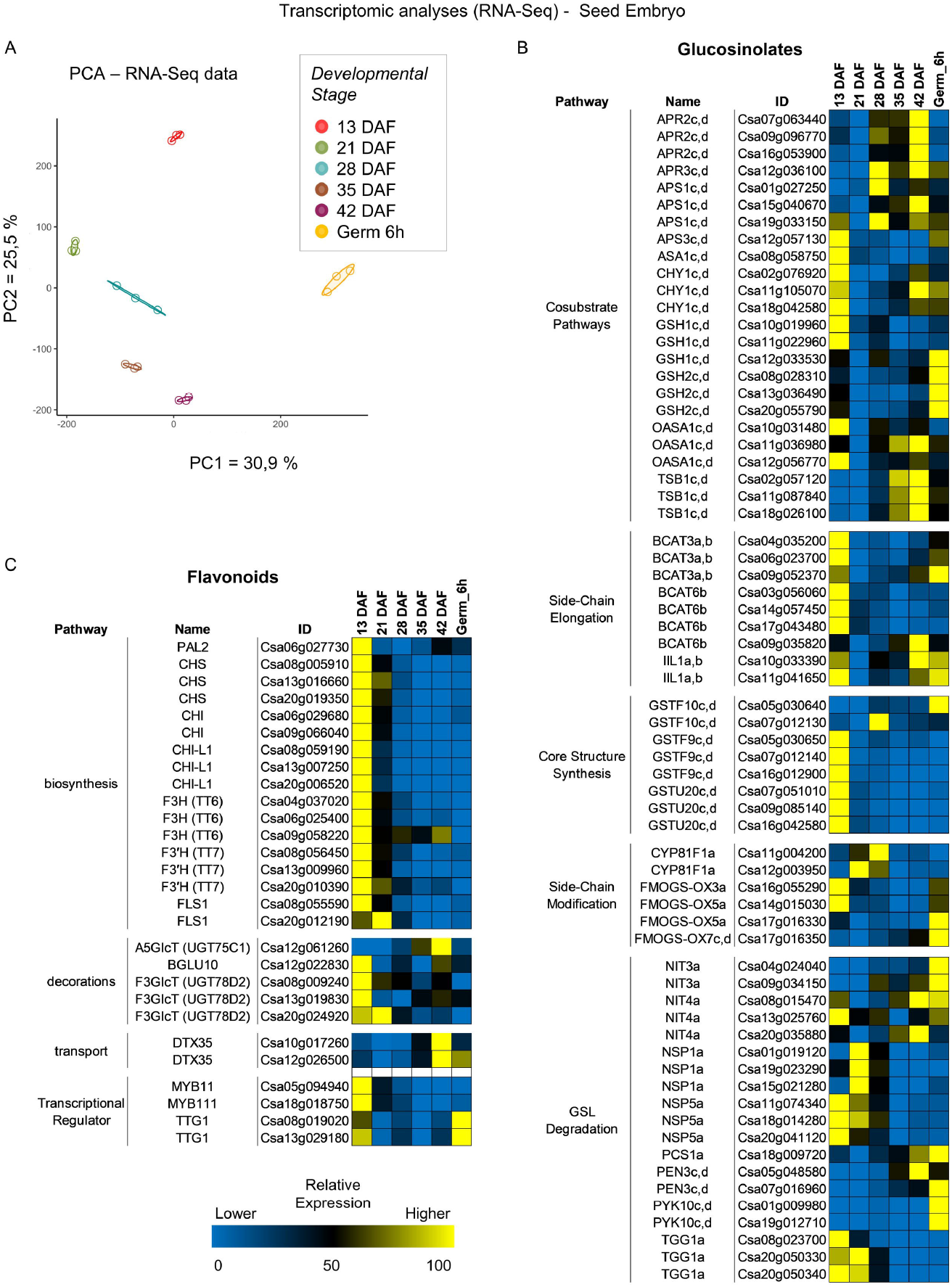
RNA-Seq transcriptomic analyses and expression of flavonoid and GSL-related genes. **A)** Principal component analyses using the RNA-Seq transcriptomic data of SCE and SE harvested at several developmental stages and during germination. DAF = Days After Flowering. **B)** Heatmap showing the expression behaviour of GSL and flavonoid –related genes during seed development in SCE and SE tissues.

## Discussion

The accumulations of SMs has a significant impact on the quality of plant-derived products. While the spatial distribution of these compounds in plant leaves and roots tissues has been described for some model and crop species (Brown et al., 2003; Czerniawski et al., 2021), very limited information and mainly restricted to primary metabolites is available about their spatial accumulation in seed tissues (Zhang et al., 2021). Given their major impact on seed quality, elucidating SM distribution in seed coat, endosperm and embryo during development and germination is of main interest in Brassicaceae species, including the oilseed crop *C. sativa* (Kagale et al., 2014; Faure and Tepfer, 2016; Zanetti et al., 2021). In the present study we combined untargeted metabolomic, proteomic and transcriptomic analyses to unveil seed SM accumulation (synthesis, transport and/or modifications/decorations) in developing seed coat and endosperm, and seed embryo. Our results emphasized a large diversity of SM in camelina seeds, as shown in a previous study (Boutet et al., 2022).

A progressive but significant evolution of the specialized metabolome can be observed during embryo development. The sample replicates of the earliest stage of seed development studied (13 DAF) and the samples replicates of the latest seed development stage (dry seeds) are the most distant; and intermediary developmental stage samples are distributed between these two samples, following a chronological order. Interestingly, the embryo specialized metabolome of 6h and 24h germinating seeds appear to be similar to those of seed embryo in the early and middle seed developmental stages, respectively (Figure 2C). Hence, these data suggest that there is a considerable specialized metabolome change/shift occurring in the embryo from dry seed to germinating seed and that the metabolome is afterwards following an evolution equivalent to the one that takes place during seed development. Similar results have been observed for the SCE, for which seeds harvested at 13 DAF and germinating seeds at 24h after imbibition showed a similar specialized metabolome.

RNA-Seq transcriptomic analysis revealed that the expression of a majority of genes involved in the biosynthetic pathways of GSLs, flavonoids, cinnamic acids, and monolignols peaks at the early seed developmental stage (13 DAF) in the embryo (Figure 10 and Figure S6). Consistently, a large part of the enzymes involved in the synthesis of GSL, flavonoid, cinnamic acid and monolignol were accumulated in the embryo between 13 DAF to 21 DAF (Figure 5B), reflecting a delay between gene expression and the accumulation of the corresponding proteins. Interestingly, SM-related enzymes from SCE were also mostly accumulated at early stage of seed development (Figure 5B; Table S3). However, SMs displayed contrasting accumulation patterns between both seed tissues (Figure 3). Indeed, whereas both SE and SCE had some groups of SMs highly accumulated at 13 DAF, a large part of SCE SMs were mainly accumulated at 35 and 42 DAF stages, whereas the majority of embryo SMs were highly accumulated at dry seed stage (Figure 3). Both of those SCE and SE SM clusters were enriched in AA and derivatives, cinnamic acids and flavonols, indicating that those metabolic categories are synthesised and/or stored in SCE during seed development and transported to SE between the maturation (42 DAF) and dry seed stage (Figure 3). Hence, those results emphasize the importance of SCE tissues, at the interface between the mother plant and the embryo, in the support of embryonic growth (Yan et al., 2014). PCA analyses on untargeted metabolomic data highlighted that, while both factors are impacting SMs diversity, metabolite signature seems more similar within the same tissue during seed development than in different tissues at a specific developmental stage (Figure 2C). For example, proanthocyanidins (PAs) are specifically accumulated in *Arabidopsis thaliana* seed coat from the early seed developmental stages (1-2 DAF) and reach the maximum peak at 6-7 DAF for Col-0 ecotype and 12 DAF for Cvi-0 ecotype (Routaboul et al., 2012; Corso et al., 2020). In addition, PA accumulation is paralleled by the expression of the *BANYULS* gene, which is involved in their synthesis (Debeaujon et al., 2003). This trend is confirmed in camelina seeds, where the anthocyanidin reductase (ANR) protein, encoded by a *BANYULS* homolog (Xie et al., 2003), showed the highest expression peak at 13 DAF in SCE samples (Figure 9A) and, consistently, the flavan-3-ols derivatives (catechin, epicatechin and procyanidin) are highly accumulated at 13, 21 and 28 DAF (Figure 9B). Differently from the aglycone forms, catechin and epicatechin glucosides (as most glycosylated and acylated flavonoids) are accumulated at later stages of seed coat development in camelina, as shown for *Vitis vinifera* seeds (Zerbib et al., 2018) (Figure 9B), consistent with an increase stability of these glycosylated metabolites, as previously shown by Raab et al. (2010).

Besides flavonoids, the defensive and antinutritional GSLs and related ITC compounds, deriving from GSL degradation, showed contrasting accumulation pattern when comparing SCE to embryo tissues, in camelina seeds (Figure 6). While some GSLs have a negative impact on seed oil and protein quality and digestibility, and are in some cases toxic for animals and humans, others are powerful antioxidants with nutritional and health benefits (Traka, 2016; Di Gioia et al., 2019). Hence, they have been a target of domestication for some Brassicaceae (e.g. Brassica and Camelina species) for which many efforts are still being done to modulate the content or nature of GSLs in seeds (Nour-Eldin et al., 2017; Hölzl et al., 2023; Mann et al., 2023). Nevertheless, GSLs are antioxidants accumulated in response to abiotic stresses (Rao et al., 2021; Mácová et al., 2022) and are essential for plant resistance to pathogens, so much so that their systematic elimination could prove counterproductive.

Our data highlighted that the different GSL spatial distribution, accumulation and, hence, functions in seeds, observed among *C. sativa*, Arabidopsis and Brassica species were mainly due to GSL side-chain length, GSL transport and potentially GSL chemical modifications (decorations). Long chain C8-C11 methylsulfinylalkyl GSLs, synthetized in camelina (C8-C11) and Arabidopsis (only C8), were indeed accumulated in the seed coat and endosperm, while mid- and short-chain C3-C7 methylsulfinylalkyl GSLs were accumulated in the embryo of Brassica and Arabidopsis species studied in this work. These differences could be due to the affinity of GSL transporters for metabolites depending on the side-chain length.

The contrasting pattern of GSL and ITC accumulation among these three Brassicaceae species could be partially due to the absence of benzoylated and sinapoylated GSLs in camelina and brassica seeds, whereas these compounds accumulate at high level in Arabidopsis embryos (Kliebenstein et al., 2007; Lee et al., 2012; Bussell et al., 2014; Sanden et al., 2023). In particular, sinapoyl- and benzoyl-GSLs could be more stable and less toxic than non-decorated GSLs and might impact their degradation to ITC and nitriles and the transport to other seed tissues and/or cellular/vacuolar compartments, thus facilitating their accumulation in the embryo. Nevertheless, no clear information is available regarding the biological activity and role of acylated-GSLs in seeds, which deserves further investigation. Regarding the distribution of ITCs, it could be suggested that long-chain ITCs (C>8) can be accumulated in the embryo rather than the SC because they may present lower toxicity levels than shorter ITCs. Indeed, some reports show that the ITC toxicity against pathogens varies depending on ITC chemical structure, including the carbon chain length (Matthäus and Zubr, 2000; Wilson et al., 2013; Abdel-Massih et al., 2023).

Interestingly, fewer SMs showed peaks of accumulation at germination stages (6h Germ and 24h Germ) compared to the seed developmental stages for both tissues (Figure 3). This suggests that many SMs are degraded rather than synthetized during germination. Similarly, there were fewer proteins strongly accumulated at germination stages compared to seed developmental stages for both tissues (Figure 5). However, it is worth mentioning that glycosyltransferases were found to be highly expressed in SE at both 6h and 24h germination stages (Figure 5). This could be linked to starch and sugar remobilization occurring during the germination and establishment of the seedling. Likewise, carbohydrates metabolic category was found to be specifically enriched upon seed germination (Figure 3). Moreover, 9-Methylsulfinylnonyl, 10-Methylsufnyldecyl and 11-Methylsulfinylundecyl ITCs were found to decrease progressively from dry seed stage to 24h Germination stage (Figure 7). This decrease of ITCs was concomitant with an increase in expression of several enzymes involved in GSL degradation, and thus potentially ITC production (Figure 10), suggesting that ITCs may have a role during camelina seed germination by providing for instance an important source of AA, sulfur atoms or other molecules.

To conclude, GSLs, flavonoids, cinnamic acids, and monolignols are largely synthetized and accumulated in camelina seeds and, therefore, determine their nutritional quality and play a key role in seed interaction with biotic and abiotic environments. Characterizing the spatial dynamics of seed SMs will contribute to the development of crops with an optimized distribution of beneficial and toxic metabolites for seeds quality and animal nutrition or to the development of new strategies (e.g. fractionation or germination) to reduce the level of undesired compounds after harvest.

## Experimental procedures

### Plant culture and seed harvest

*Camelina sativa* (L.) “Céline” genotype was used for the plant culture and analyses. Seeds of the three biological replicates (10 plants/replicate) were sown with a gap of two weeks between each replicate and were cultivated in different part of the greenhouse. Plants were cultivated in 16 cm diameter pots (from TEKU) in a greenhouse: 16h light (23-25 °C) / 8h dark (15°C) (Osram Powerstar E40 HQI-BT 400W lights). Relative humidity room was set on 45%. A nutrient solution containing 140 milligrams of nitrogen per liter (140 mg N L⁻¹) was used to irrigate the plants.

Flowers were tagged just before opening to count opened flowers and harvest seeds at defined stages (Sakai et al., 2018). Seed embryos and coats/endosperms were separated and harvested at 13, 21, 28, 35, 42 DAF, at dry mature seed stage, and at 6 hours and 24 hours, after imbibition, during germination.

For seed pictures acquisition, siliques from each time-point were carefully dissected to harvest the seeds. Chloral hydrate solution (chloral hydrate:H_2_O_2_:glycerol (8:2:1, w/v/v) was used to bleach seed coats and fix the samples that were placed between slide and slip (Boisson et al., 2001). Seeds were then stored for 24h to 72h at 4°C before imaging. Images were acquired on Axioplan 2 imaging epifluorescence microscope after several weeks of seed coat clearing (Zeiss, Göttingen, Germany).

*Arabidopsis thaliana* “Columbia-0” (Col-0) genotype was used in this study. Seeds were sown directly in fertilized soil (3 biological replicates, three plants/replicate). Plants were cultivated in 7.5 x 7.5 cm pots in a growth chamber with long day photoperiod: 16h light (21°C) - 8h dark (19°C). Dry seeds were harvested and kept at -20°C until dissection.

For *Brassica napus* species, “Aviso” and “Major” genotype, showing contrasting glucosinolate accumulation, were used. Seeds were sown directly in fertilized soil. After 4 weeks the plants were transferred in a vernalisation room for 10 weeks (7,5 x 7,5 cm pots, at 4°C, with a 16/8h day/night photoperiod). Next, plants were transferred in 19L pots and cultivated during 5 months in a growth chamber with long day photoperiod: 14h of light (21°C) and 10h dark (13°C). Plants were watered 2 times a day with a nutritional solution (140mg of nitrogen / liter). Dry seeds were harvested and kept at -20°C until the dissection.

## Specialized metabolites extraction and analysis

### Metabolite extraction and injection

Untargeted metabolomic data analyses and date were described in (Boutet et al., 2022). Briefly, metabolites were extracted from 20 mg of seed coats and embryos using a MeOH: Methyl-tert-butyl: H2O (1:3:1) buffer, while polar and semipolar metabolites were separated from the oil and protein fractions using a MeOH: H2O (1:3) buffer. Polar and semipolar metabolite fraction was used for metabolomic analyses. Untargeted metabolomic data were acquired using a UHPLC system (Ultimate 3000 Thermo) coupled to quadrupole time of flight mass spectrometer (Q-Tof Impact II Bruker Daltonics, Bremen, Germany). For chromatographic separation a Nucleoshell RP 18 plus reversed-phase column was used. Samples were injected in positive and negative ionisation mode (ESI+ and ESI-).

### Data processing and ion annotation

ESI+ and ESI-were processed using MZmine 2.52 software (http://mzmine.github.io/). Metabolite annotation was performed in four steps: i) LC-MS/MS data were compared with a home-made library (IJPB chemistry and metabolomic platform) containing more than 400 standards or experimental common features; ii) LC-MS/MS data were also searched against the available MS2 spectral libraries (Massbank NA, GNPS Public Spectral Library, NIST14 Tandem, NIH Natural Product and MS-Dial); iii) a molecular network analysis was used to assign not-annotated metabolites to a chemical family (see Boutet *et al*., 2022 and Olivon *et al*., 2018 for further information); iv) Sirius software (https://bio.informatik.uni-jena.de/software/sirius/) was used to assign a putative annotation to metabolic features that were not annotated during the previous steps. Raw data were normalised on the internal standard (Apigenin) and weight of seeds used for the extraction.

### Statistical analyses of metabolomic data

Statistical analyses were performed using Metaboanalyst 5.0 software (Pang et al., 2021; Pang et al., 2022). In particular, an ANOVA has been conducted to identify differentially accumulated metabolites among seed developmental stages, separately for SCE and SE (adjusted p value < 0.05).

## Transcriptomic analyses (RNA-Seq)

### RNA extraction

RNAs were extracted with the PicoPure™ RNA Isolation Kit (ThermoFisher) following the corresponding protocol with slight modifications. Briefly, 2 to 5 mg of seeds were grinded with some polyvinylpolypyrrolidone (PVPP; phenolic compound absorbing agent) in liquid nitrogen. 100 µL of extraction buffer was added to each sample. Following, samples were vortexed, incubated at 42°C for 30 min (500 rpm) using a ThermoMixer^TM^ C (Eppendorf), and centrifuged (2 min, 3000 g). Supernatants were collected, without picking-up the pelleted material, and placed in the pre-conditioned RNA purification columns to be centrifuged (1 min, 16 000 g). The cell extracts obtained were mixed with 100 µL of 70 % ethanol by pipetting up and down. The mixtures were placed into the pre-conditioned purification columns which were centrifuged 2 min at 100 g, to allow RNA binding to the columns, immediately followed by a centrifugation to remove flow-through (30 sec, 16 000 g). The columns were washed by adding 100 µL of wash buffer 1 and centrifugation (1 min, 8 000 g). DNase treatment was made by adding 40 µL of DNase solution mix (10μl DNase I Stock solution + 30μl Buffer RDD; RNase-Free DNase Set, Qiagen) to each column, followed by 15 min of incubation (ambient temperature) and washing of the column (addition of 40 µL of wash buffer 1 and centrifugation 30 sec at 8 000 g). Columns were washed twice with 100 µL of wash buffer 2 (2 min at 8 000 g) and a last centrifugation was made after flow-through waste discard to remove all traces of wash buffer (1 min, 16 000 g). 12 µL of elution buffer were added to each column, which were incubated 1 min at ambient temperature before RNA elution centrifugations (1 min at 1 000 g followed by 1 min at 16 000 g). Obtained RNAs were stored at -80 °C until sequencing.

### RNA sequencing

Sequencing of messenger RNAs (mRNAs) was performed by BGI Genomics (Hong Kong, https://www.bgi.com). Prior to cDNA library construction (Eukaryotic Strand-specific Transcriptome), quality and concentrations of mRNAs were checked again with the bioanalyzer from BGI platform. The sequencing (paired-end reads of 150bp, 70M clean reads per sample) was performed with DNA Nanoballs Technology (DNBSEQ^TM^). After sequencing, raw reads were filtered by the BGI platform with the filtering software SOAPnuke (version 1.5.2, https://github.com/BGI-flexlab/SOAPnuke) (Cock et al., 2009; Chen et al., 2018) (Parameters used: l 15 -q 0.2 -n 0.05) to reach a Phred+33 fastq quality score.

### Annotation and statistical analysis of transcriptomic data

High quality raw reads (Céline strain) were aligned against the *Camelina sativa* genome (strain: DH55, genome assembly: GCA_000633955.1, annotation: https://www.camelinadb.ca/downloads/:_gff_and_fasta_to_be_made_available_if_no_more_available_in_this_site) using GSNAP (gmap-2021-12-17) with a set of options allowing to find novel splicing and to be tolerant to SNP between DH55 and Céline (VCF files, to be made available). htseq-count (HTSeq-0.12.4-python2) was used with the following options (-m intersection-nonempty--secondary-alignments=ignore -a 20 -s no) to count the number of reads by gene. Orthologous genes of *Arabidopsis thaliana* were searched in *Camelina sativa* using Plant.ensembl website (using Biomart) and CamRegBase (synthelogs data).

Statistical analyses were performed to identify Differentially Expressed Genes using the Edge R package (Robinson et al., 2010). Statistical analyses were performed to identify genes that were differentially expressed among developmental and germination stages in the SCE and SE separately (FDR < 0.05).

## Proteomic analyses (DIA)

### Sample preparation and analysis

Five µg of each protein extract were prepared using a modified Gel-aided Sample Preparation protocol (https://www.ncbi.nlm.nih.gov/pmc/articles/PMC4409837/). Samples were digested with trypsin/Lys-C overnight at 37°C. For nano-LC fragmentation, protein or peptide samples were first desalted and concentrated onto a µC18 Omix (Agilent) before analysis.

The chromatography step was performed on a NanoElute (Bruker Daltonics) ultra-high-pressure nano flow chromatography system. Approximatively 200ng of each peptide sample were concentrated onto a C18 pepmap 100 (5mm x 300µm i.d.) precolumn (Thermo Scientific) and separated at 50°C onto a reversed phase Reprosil column (25cm x 75μm i.d.) packed with 1.6μm C18 coated porous silica beads (Ionopticks). Mobile phases consisted of 0.1% formic acid, 99.9% water (v/v) (A) and 0.1% formic acid in 99.9% ACN (v/v) (B). The nanoflow rate was set at 250 nl/min, and the gradient profile was as follows: from 2 to 30% B within 70 min, followed by an increase to 37% B within 5 min and further to 85% within 5 min and reequilibration.

MS experiments were carried out using a TIMS-TOF pro mass spectrometer (Bruker Daltonics) with a modified nano electrospray ion source (CaptiveSpray, Bruker Daltonics). A 1400 spray voltage with a capillary temperature of 180°C was typically employed for ionizing. MS spectra were acquired in the positive mode in the mass range from 100 to 1700 m/z and 0.60 to 1.60 1/k0 window. In the experiments described here, the mass spectrometer was operated in PASEF DIA mode with exclusion of single charged peptides. The DIA acquisition scheme consisted of 16 variable windows ranging from 400 to 1200 m/z.

### Protein identification

Database searching and LFQ quantification (using XIC) was performed using DIA-NN (version 1.6.1) (Demichev et al., 2020). The UniProt *Camelina sativa* database was used for library-free search / library generation. For RT prediction and extraction mass accuracy, we used the default parameter 0.0, which means DIA-NN performed automatic mass and RT correction. Top six fragments (ranked by their library intensities) were used for peptide identification and quantification. The FDR was set to 1% at the peptide precursor level. The variable modifications allowed were as follows: Nterm-acetylation and Oxidation (M). In addition, C-Propionoamide was set as fix modification. “Trypsin/P” was selected. Data were filtering according to a FDR of 1%. Cross-run normalisation was performed using RT-dependent.

### Identification of differentially expressed proteins

To quantify the relative levels of protein abundance between different groups, datas from DIA-NN were then analysed using DEP package from R. Briefly, proteins that are identified in 2 out of 3 replicates of at least one condition were filtered, missing data were imputed using random draws from a manually defined left-shifted Gaussian distribution and differential enrichment analysis was based on linear models and empherical Bayes statistic. A 1.2-fold increase in relative abundance and a 0.01 pvalue were used to determine enriched proteins. ANOVA were performed from PERSEUS to determine those enriched proteins. A 1.2-fold increase in relative abundance and a 0.05 FDR were used.

### Glucosinolate distribution in Brassicaceae seeds

Seed embryos and coats/endosperms were separated from dry seeds of *Arabidopsis thaliana* (L.) “Columbia-0” (Col-0) ecotype and *Brassica napus* (L.) “Major” and “Aviso” genotypes.

Specialized metabolite extraction and metabolomic analyses were performed as presented previously (see “Specialized metabolites extraction and analysis”). 200 seeds and 25 mg were extracted for Arabidopsis and Brassica samples respectively.

Arabidopsis LC-MS/MS data were first normalized according to the seed tissue proportion (Seed coat and endosperm: 10 % and Seed embryo: 90 %), following data were normalized by the total sum of accumulation per sample in each mode (ESI + and ESI-), multiplied by 1000.

### Clustering analyses

The R package ‘pheatmap” was used to perform hierarchical clustering analysis (hclust) and create heatmaps using omics data.

## Acknowledgements

This work was supported by AgroParisTech (grant “Appel à projets scientifiques 2020” to M.C. and L. R.), the “Département de Biologie et Amélioration des Plantes” (BAP) of INRAE (grant “Appel à projets scientifiques BAP 2020” to M.C.), by the project ‘Seed MetSpe’ of the Labex Saclay Plant Sciences-SPS (ANR-10-LABX-0040-SPS to L.L. and M.C.) and by a project financed by Plant2Pro Carnot institute (‘BrassiMet’ to M.C.).

The IJPB benefits from the support of Saclay Plant Sciences-SPS (ANR-17-EUR-0007). This work has benefited from the support of IJPB’s Plant Observatory platforms PO-Chem and PO-Plants. We are especially thankful to Patrick Grillot (IJPB, INRAE-Versailles) for his help during the plant culture in the greenhouse.

The authors would like to thank as well Dr Mark Tepfer (retired, former IJPB, INRAE) for the constructive discussion on *Camelina sativa* species.

## Data availability

The metabolomic data and metadata on *Camelina sativa* have been deposited at the MassiVE data repository portal with the identifier MSV000094688 (doi:10.25345/C5HH6CH59).

The transcriptomic data have been deposited at the National Center for Biotechnology Information (NCBI) Transcriptome Shotgun Assembly Sequence Database (TSA) with BioProject identification PRJNA835304.

The proteomic data have been deposited at the iProX (integrated Proteome resources) data repository portal with the identifier IPX0008673001

The metabolomic data and metadata on *Arabidopsis thaliana* have been deposited at the MassiVE data repository portal with the identifier MSV000095844 (doi:10.25345/C51J97K84).

The metabolomic data and metadata on *Brassica napus* have been deposited at the MassiVE data repository portal with the identifier MSV000095843 (doi:10.25345/C5599ZD2H).

## Author contributions

MC and LR directed and designed the research. MC, DDV, TF, DG and CB grew the plants and harvested seed coat and embryos at defined developmental stages. SB and JCT generated the LC-MS/MS data and performed the initial quality analyses on the metabolomic data. SB, FP, LB, NK, CB and JCT performed the metabolites annotation with MetGem, SIRIUS and using MS/MS spectra. MC, LB, CB, SB, FP, NK and LL analysed and interpret the untargeted metabolomic data. BC prepared the samples for proteomic analyses. BB performed the proteomic analyses. MC analysed transcriptomic data. MC, LB and CB performed the statistical analyses on multiomic data. MC, LB and CB generated all Fig.s and supplemental material. MC and LB wrote the paper, which was edited by CB, LR and LL. All the authors commented on and approved the manuscript.

