## Supplementary figures and images for "Multi-omic analyses unveil contrasting composition and spatial distribution of specialized metabolites in seeds of *Camelina sativa* and other Brassicaceae"

### Fig. S1

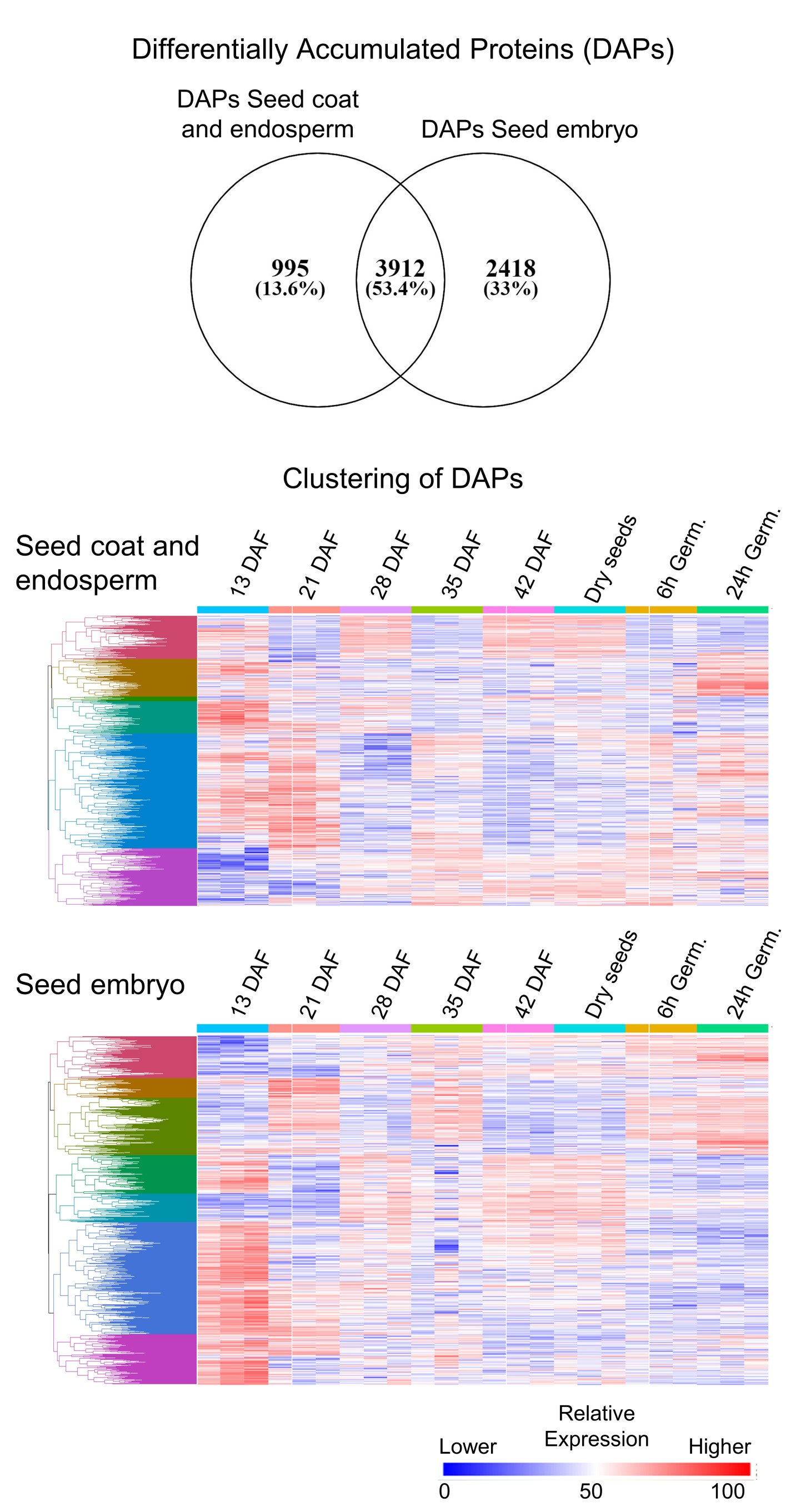

### Fig. S2

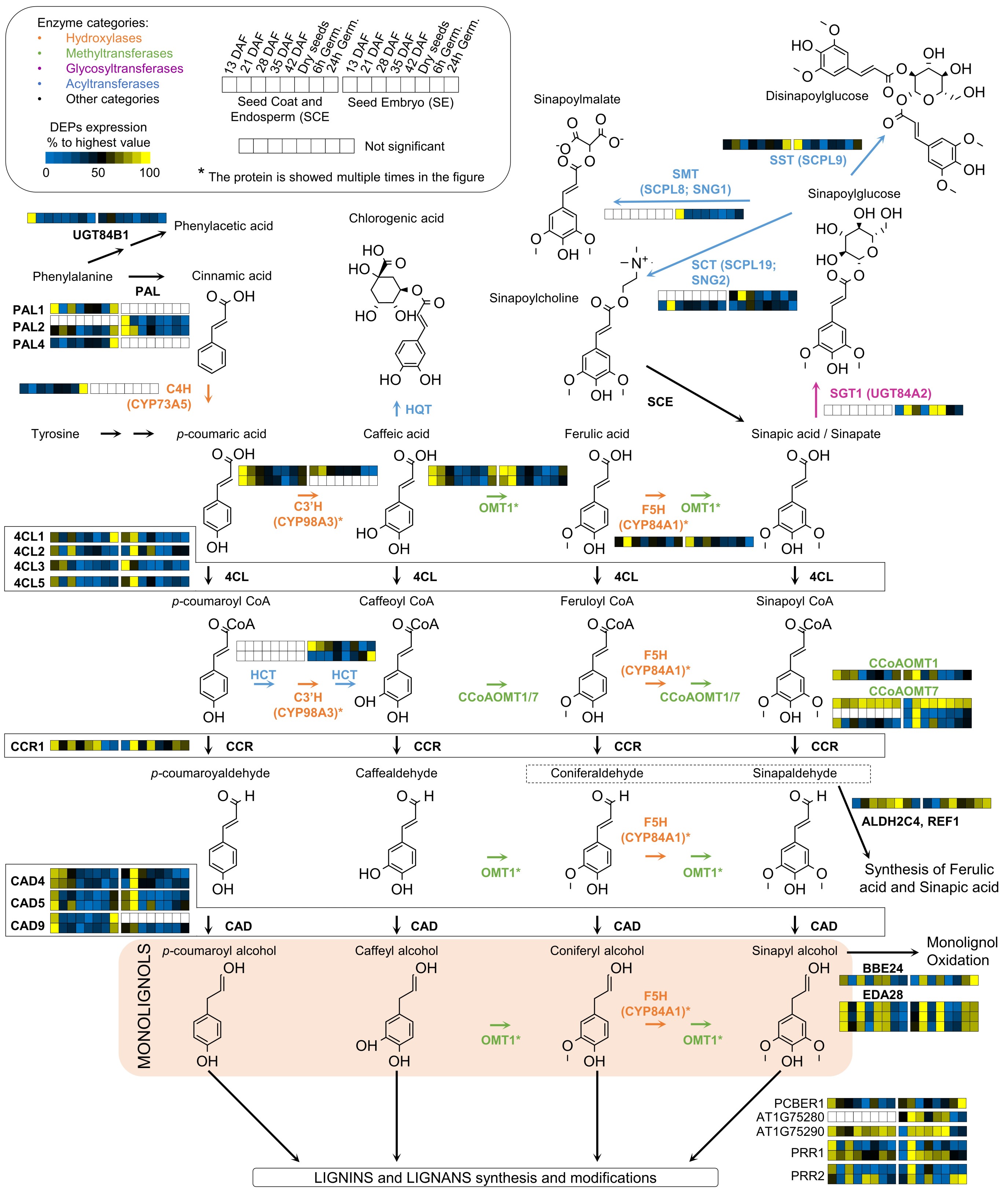

### Fig. S3

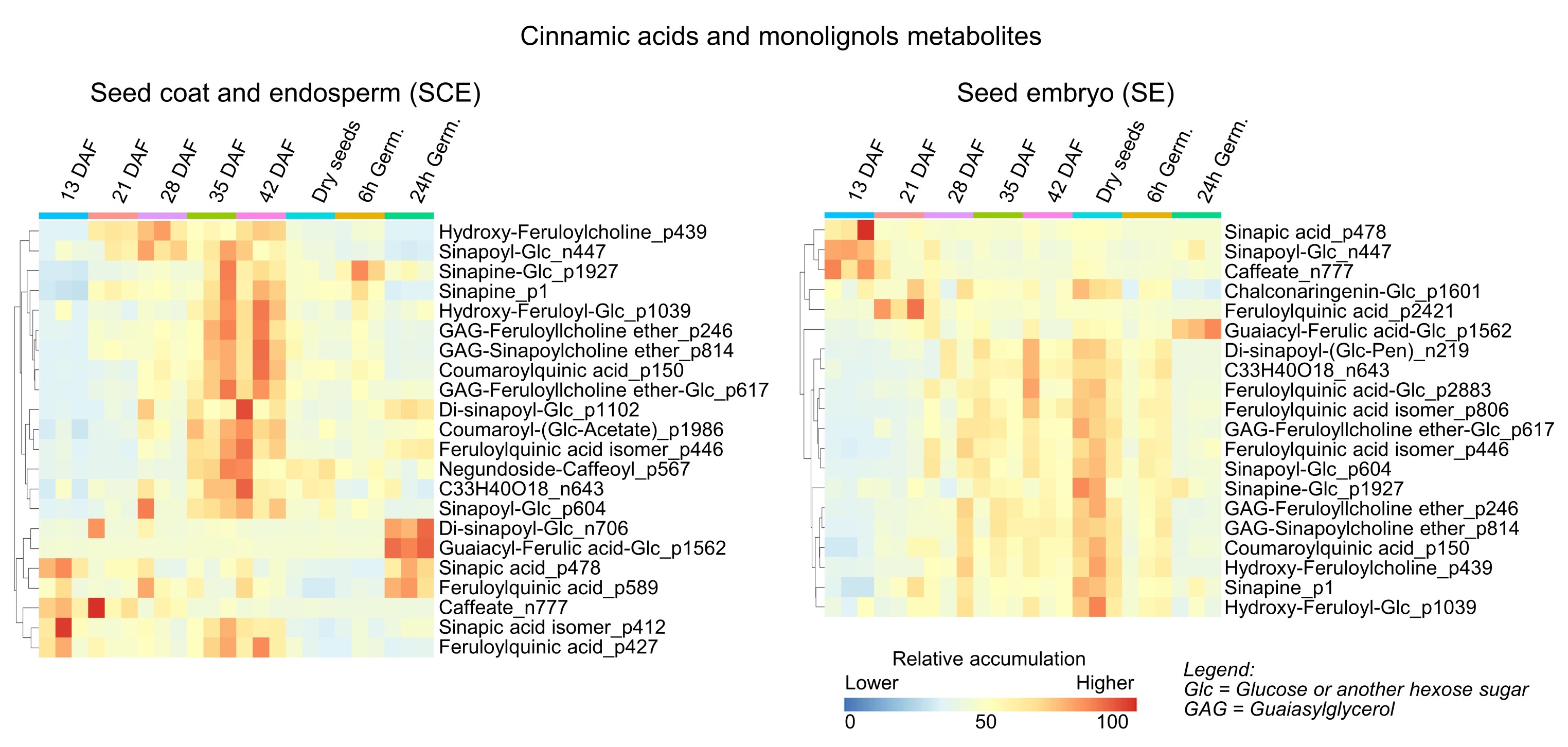

### Fig. S4

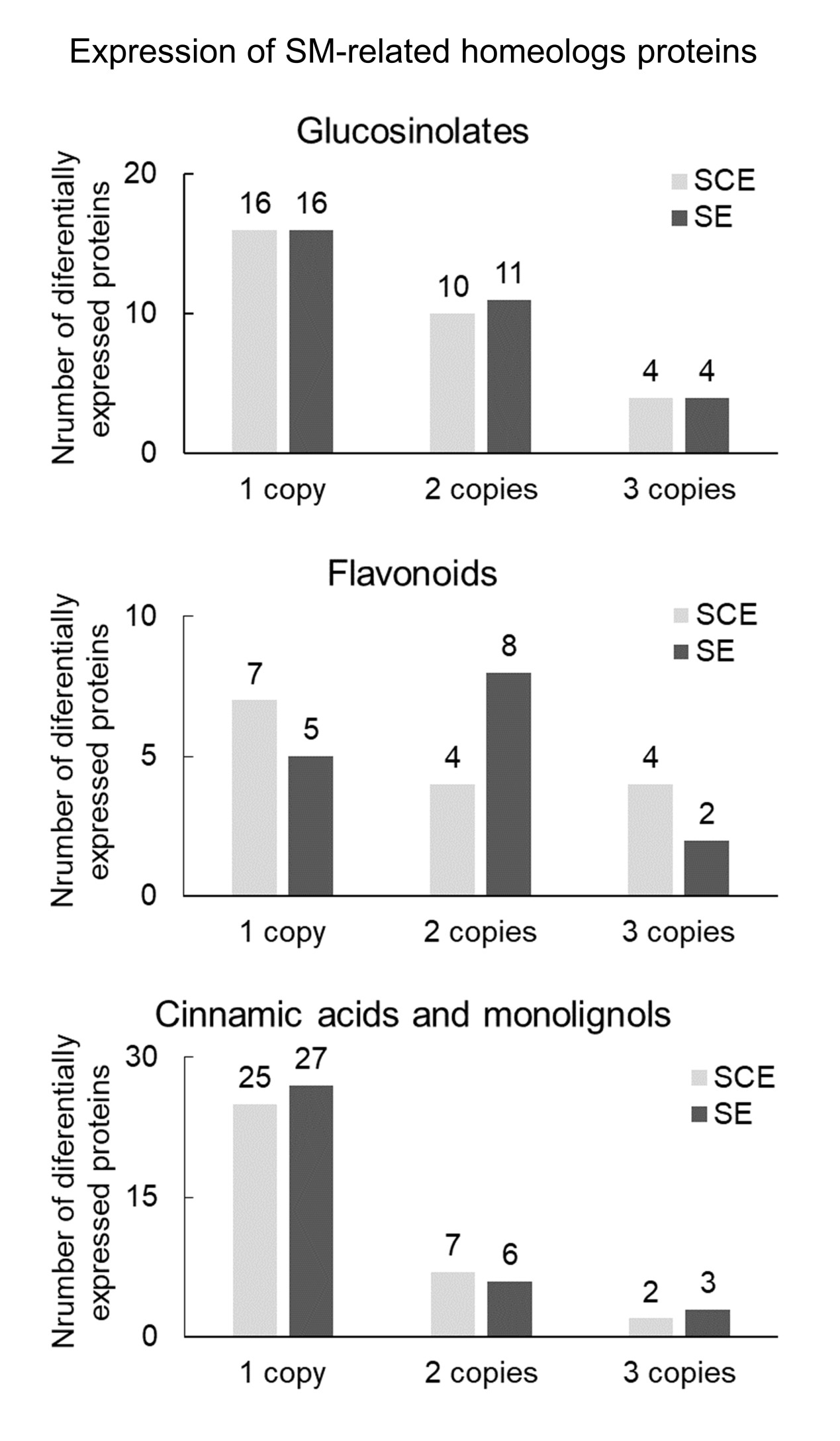

### Fig. S5

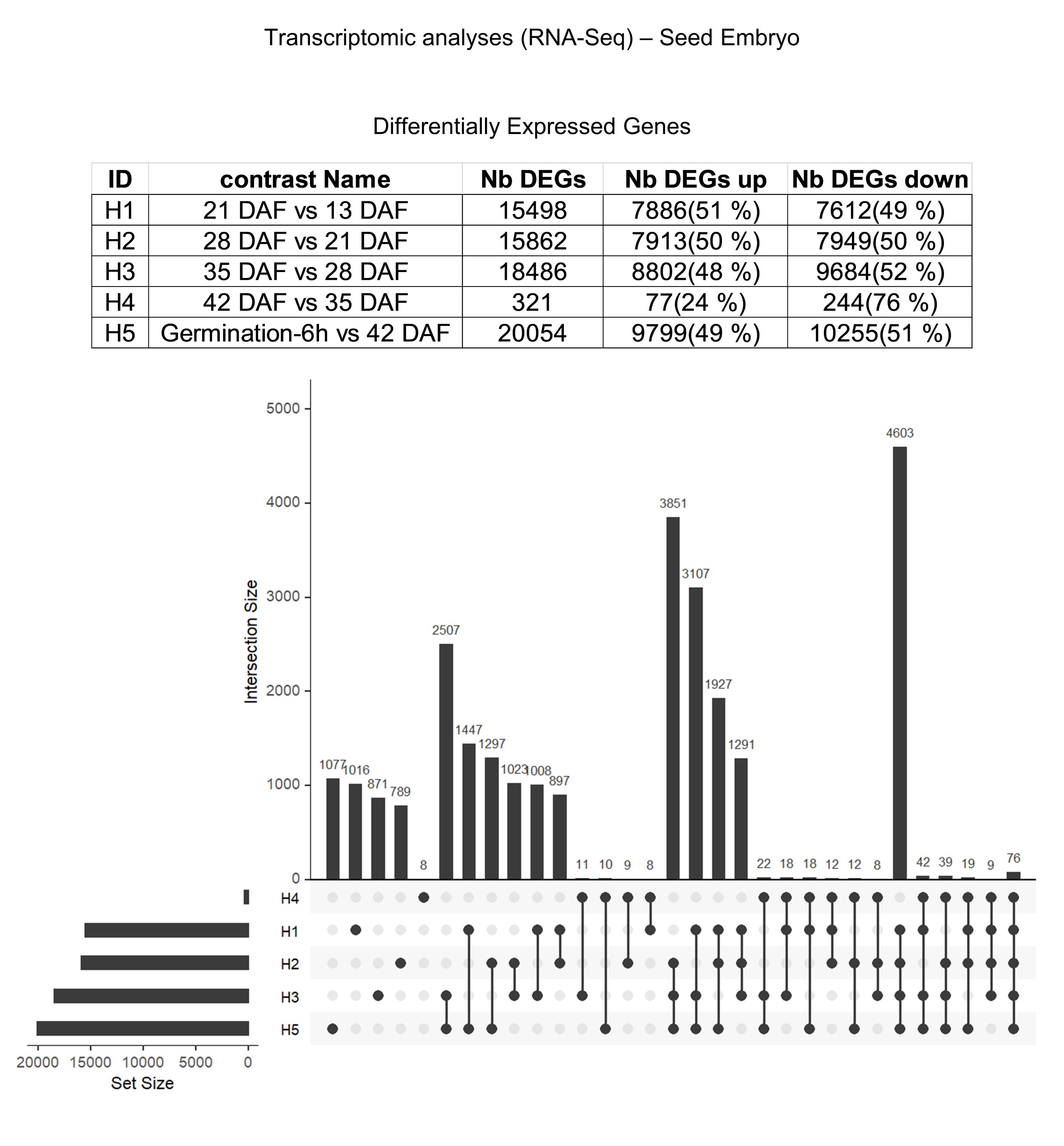

### Fig. S6

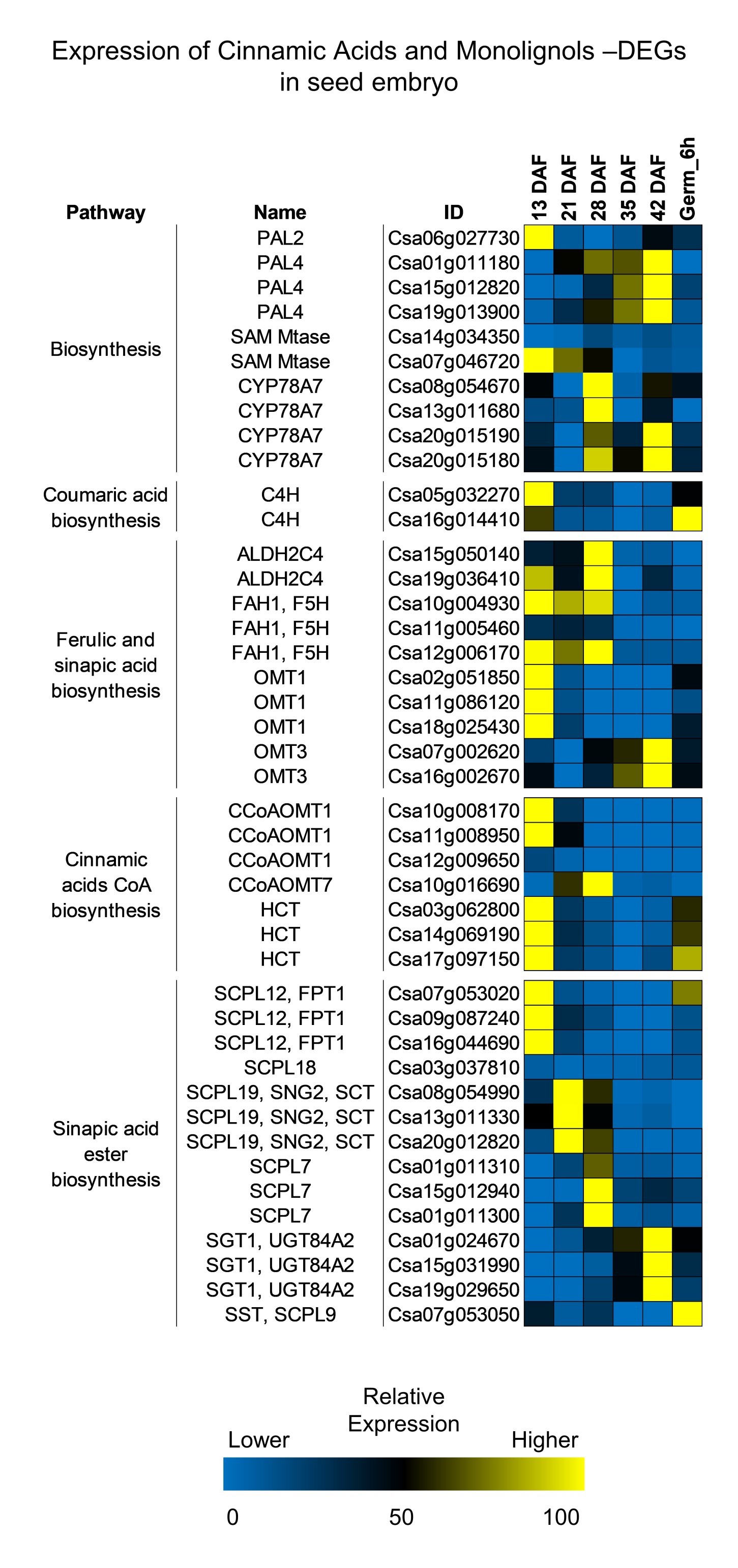
