## Supplementary material for "Multi-omic analyses unveil contrasting composition and spatial distribution of specialized metabolites in seeds of *Camelina sativa* and other Brassicaceae": Table S1

**Table S1.** Top-10 most intense specialized metabolites in camelina seed coats (a) and embryos (b).

a) Seed coat and endosperm specialized metabolites:

| *ID* | *Annotation* | *Category* | *Average Intensity* |
| --- | --- | --- | --- |
| p1 | Sinapine | Cinnamic acids/alkaloids | 118 |
| p2 | 10-Methylsulfinyldecyl isothiocyanate | Isothiocyanates | 86 |
| p4 | 9-Methylsulfinylnonnyl isothiocyanate | Isothiocyanates | 43 |
| p5 | Rutin (quercetin-3-Glucoside-Rhamnoside) | Flavonols | 42 |
| p8 |  | Cinnamic acids | 36 |
| p9 | 11-Methylsulfinylundecyl isothiocyanate | Isothiocyanates | 29 |
| p7 | Adenosine | Nucleosides and derivatives | 25 |
| n2 | Glucoarabin (9-methylsulfinylnonyl GLS) | Glucosinolates | 25 |
| p298 | Epicatechin | Flavan-3-ols and PAs | 24 |
| p2553 | C_17_H_32_NO_7_S_2_^+^ | Glucosinolate derivative | 19 |

b) Seed embryo specialized metabolites:

| *ID* | *Annotation* | *Category* | *Average Intensity* |
| --- | --- | --- | --- |
| p1 | Sinapine | Cinnamic acids/alkaloids | 71 |
| p5 | Rutin (quercetin-3-Glucoside-Rhamnoside) | Flavonols | 64 |
| p13 | C_11_H_2_0N_2_O_5_S | Aa and derivatives | 62 |
| p18 | Quercetin-(Xyloside-Rhamnoside-Glucoside) | Flavonols | 62 |
| p2 | 10-Methylsulfinyldecyl isothiocyanate | Isothiocyanates | 53 |
| p4 | 9-Methylsulfinylnonnyl isothiocyanate | Isothiocyanates | 48 |
| p29 | C_13_H_25_0NOS | Sulfoxides | 45 |
| p51 | (2S)-2-amino-5-[[(1S)-1-carboxy-2-methylsulfanylethyl]amino]-5-oxopentanoic acid | Aa and derivatives | 38 |
| p48 | gamma-Glu-Leu | Aa and derivatives | 37 |
| p7 | Adenosine | Nucleosides and derivatives | 28 |
